# Linking post-stress brain connectivity to acute cortisol reactivity using network-based inference and prediction

**DOI:** 10.64898/2026.08.14.744860

**Authors:** Emin Serin, Elvan Emurla, Christoph Bärtl, Marina Giglberger, Julian Konzok, Hannah L. Peter, Ludwig Kreuzpointner, Brigitte M. Kudielka, Stefan Wüst, Susanne Erk, Henrik Walter, Gina-Isabelle Henze

## Abstract

**Background:** Acute cortisol responses to psychosocial stress vary substantially across individuals, yet how this variability is reflected in post-stress resting-state functional connectivity (rsFC) remains unclear. Although prior work has linked stress-related endocrine responses to brain connectivity, studies have been limited by small samples, region-of-interest approaches, or a sole focus on group-level analyses. Here, we investigated whether acute cortisol increase is associated with, and can be predicted from, whole-brain post-stress rsFC.

**Methods:** We analyzed 339 healthy participants from two Scan*STRESS* datasets using complementary inferential and predictive approaches. First, we used the Network-Based Statistic (NBS) to identify connected rsFC networks associated with acute cortisol increase, controlling for age, site, and sex/hormonal status. Second, we predicted participants’ acute cortisol increase from their connectivity patterns using NBS-Predict and Connectome-Based Predictive Modeling (CPM). Together, we examined the cortisol-rsFC relationship at the population and individual levels.

**Results:** Greater cortisol responses were associated with lower post-stress rsFC within a significant distributed network comprising 258 connections among 78 regions, centered on thalamic nuclei and pallidal regions and extending to default-mode, limbic, orbitofrontal, and cerebellar regions. Sex-stratified analyses revealed a significant negative association only in females, but formal sex-difference contrasts were not significant. NBS-Predict and CPM yielded modest but significant out-of-sample prediction, with predictive networks converging on subcortical and posterior cingulate regions.

**Conclusions:** Post-stress rsFC carries convergent inferential and predictive information about individual HPA-axis reactivity. Stronger cortisol responses were characterized by reduced connectivity within a distributed subcortical-cingulate network, supporting a network-level perspective on neural-endocrine coupling following acute stress.

## INTRODUCTION

Acute and chronic stress are implicated in the development of high-burden conditions, including cardiovascular disease and mental disorders such as depression and anxiety (Daviu et al., 2019; Kivimäki & Steptoe, 2018; Yang et al., 2015). Stress responses vary substantially across individuals and are regulated in part by the hypothalamic-pituitary-adrenal (HPA) axis. Cortisol secretion represents a central adaptive component of HPA-axis activity, enabling the organism to respond to internal and external demands; however, sustained or dysregulated cortisol responses may contribute to stress-related health problems. Elucidating the mechanisms underlying interindividual variability in cortisol responses may therefore help identify profiles of susceptibility and resilience to stress-related disorders (Zänkert et al., 2019).

A key candidate mechanism is the organization of large-scale brain networks. Acute stress and HPA-axis engagement have been associated with changes in brain activity and connectivity (Hermans et al., 2011, 2014), particularly within the Salience Network (SN), Default Mode Network (DMN), and Central Executive Network (CEN) (Van Oort et al., 2017), which are also implicated in a range of neurological and psychiatric conditions (Menon, 2011). Although prior task-based functional magnetic resonance imaging (fMRI) studies using psychosocial stress paradigms such as Scan*STRESS* and the Montreal Imaging Stress Task (MIST) have yielded heterogeneous findings, a recent harmonized Scan*STRESS* mega-analysis identified robust stress-related activation and task-based connectivity patterns across large-scale networks and linked these responses to acute cortisol increases (Henze et al., 2025). Notably, the same study further revealed sex- and age-related variability, underscoring interindividual heterogeneity in neural stress processing.

While task-based studies provide important insight into transient, task-evoked neural responses during acute stress exposure, they do not fully capture how stress and HPA-axis engagement are reflected in the brain’s intrinsic functional architecture after the stressor. Resting-state functional connectivity (rsFC) measures the intrinsic organization of large-scale brain networks in the absence of explicit task demands (Biswal et al., 1995) and is sensitive to stress-related changes in network-level organization that may persist beyond the task period (Hermans et al., 2011; Veer et al., 2011). Importantly, post-stress rsFC should not be interpreted as a neutral baseline or purely trait-like resting state measure. Acute cortisol responses unfold with a characteristic temporal delay, typically reaching peak levels after stressor onset rather than during the immediate task period (Herman et al., 2016; van Marle et al., 2010). In the present study, resting-state data were acquired immediately following psychosocial stress induction and therefore fell within the temporal window in which cortisol levels are expected to be elevated. Post-stress rsFC can thus be conceptualized as a perturbed network state sampled during ongoing endocrine stress regulation rather than as an unstressed resting condition (Clemens et al., 2017; Maron-Katz et al., 2016; Vaisvaser et al., 2013). This temporal feature motivated the present study’s focus on the increase in cortisol as the primary outcome of interest. Psychosocial stress responses are multidimensional, including endocrine, autonomic, affective, and behavioral components (Dickerson & Kemeny, 2004; McEwen, 2007). However, cortisol increase provides a well-established marker of endocrine stress reactivity (Dickerson & Kemeny, 2004; Kudielka et al., 2009) and is particularly well suited to the present design as it indexes HPA-axis reactivity during the post-stress period in which rsFC was assessed. Accordingly, the present analyses were designed to specifically characterize neural-cortisol coupling, rather than to model stress reactivity as a broader multidimensional phenotype.

Converging evidence suggests that acute stress exposure is associated with distributed changes in rsFC rather than alterations confined to a single region or network. Across studies, stress-related rsFC alterations have most consistently implicated limbic and paralimbic circuitry and its interactions with salience, default-mode, fronto-subcortical, and thalamo-cortical systems (Dimitrov et al., 2018; Ginty et al., 2019; Reinelt et al., 2019). For example, acute stress has been linked to altered amygdala coupling with cortical midline regions, including the posterior cingulate cortex (PCC) and medial prefrontal cortex (mPFC), as well as with salience-related regions such as dorsal anterior cingulate cortex (dACC), and with hippocampal regions (Vaisvaser et al., 2013; van Marle et al., 2010; Veer et al., 2011; Y. Wang et al., 2018). Other studies have reported broader reconfigurations of thalamo-cortical, fronto-as well as parieto-limbic, and large-scale network organization (Dark et al., 2020; Maron-Katz et al., 2016; Reinelt et al., 2019), including dynamic changes in connectivity over time (Kühnel et al., 2022; Quaedflieg et al., 2015). Emerging evidence further indicates that these post-stress connectivity patterns, particularly within the SN and DMN, may relate to acute cortisol responses (Luo et al., 2026; W. Zhang et al., 2019, 2020; Zhu et al., 2020) and may vary as a function of sex, hormonal status, age and personality (Dimitrov et al., 2018; Kogler et al., 2016; Mikneviciute et al., 2023; Nasseri et al., 2020).

However, the existing literature has several important limitations. Prior studies have often relied on relatively small samples, focused on selected regions or networks, and included demographically restricted or single-sex cohorts, limiting both anatomical scope and robustness. Moreover, most previous studies have used traditional univariate statistical approaches, leaving the multivariate nature of whole-brain rsFC, individual-level prediction, and out-of-sample generalizability underexplored. To date, only one study has applied multivariate machine-learning (ML) methods to distinguish pre- and post-stress rsFC and relate the resulting classification probabilities of connectivity changes (i.e., odds ratio) to acute cortisol increase (W. Zhang et al., 2020). Although this represented an important step toward individual-level multivariate analysis of stress-related rsFC, the direct prediction of acute cortisol increases from whole-brain post-stress rsFC, while accounting for potential confounders such as sex, hormonal status, and age, remains largely unexplored.

In the present study, we examined how acute cortisol increase following psychosocial stress is reflected in post-stress functional brain architecture using two complementary, large-scale, data-driven approaches: inference and prediction. In the inferential analysis, we applied the Network-based Statistic (NBS; Zalesky et al., 2010) to identify connected networks of post-stress resting-state connections associated with acute cortisol increase. In the predictive analysis, we predicted individual cortisol increase from post-stress rsFC using ML-based NBS-Predict (Serin et al., 2021) and Connectome-based Predictive Modeling (CPM; Shen et al., 2017). Together, these approaches address distinct but complementary aspects of neural-endocrine coupling: NBS tests whether cortisol increase is associated with connected network components within the sample, whereas predictive modeling evaluates whether post-stress rsFC contains information that generalizes to unseen individuals.

## METHODS AND MATERIALS

### Sample

The present study is based on a subsample of a previously published task-based mega-analysis of psychosocial stress (Henze et al., 2025). In contrast to the original mega-analysis (*N*=459, 222 females, from eight studies across four sites), the present analyses included only datasets with post-stress resting-state fMRI acquired immediately after Scan*STRESS* and available cortisol data. The final sample comprised 339 healthy participants from two sites, Berlin and Regensburg, aged 18-62 years. The Berlin sample (*M*=28.04, *SD*=6.65) was drawn from a single study (Dahm et al., 2017), whereas the Regensburg sample (*M*=26.24, *SD*=9.01) comprised data from four independent studies sharing the same Scan*STRESS* protocol, scanner, cortisol assay, and core inclusion criteria (Bärtl et al., 2024; Giglberger et al., 2023; Henze et al., 2020; Konzok et al., 2021). Consistent with the previous task-based mega analysis, only healthy participants from the Regensburg Burnout-Project were included, whereas participants with subclinical burnout symptoms were excluded (Bärtl et al., 2024; Henze et al., 2025). An additional Berlin dataset was not included because resting-state fMRI images showed incomplete coverage of superior brain regions (Noack et al., 2019).

Of the 339 participants, 146 were male (age range: 19-60 years, *M*=27.14, *SD*=8.04) and 193 were female (age range: 18-62 years, *M*=26.29, *SD*=8.91). Among female participants, 102 were tested during the luteal phase of the menstrual cycle, 80 were using hormonal contraceptives, and 11 were post-menopausal. Sample characteristics are reported in more detail in Supplementary Tables 1 and 2. The sample inclusion and exclusion criteria are also detailed in the Supplementary Methods. All participants provided informed consent, received monetary compensation, and were tested in studies approved by the respective local ethics committees.

### Scan*STRESS* paradigm and resting-state fMRI acquisition

Acute psychosocial stress was induced inside the MRI scanner using the Scan*STRESS* paradigm (Streit et al., 2014), which consisted of alternating stress and control blocks across two runs. During stress blocks, participants performed challenging arithmetic and mental-rotation tasks under time pressure, negative performance feedback, and social-evaluative observation; during control blocks, they completed simpler matching tasks. The Regensburg studies used an optimized version of the original protocol to enhance cortisol responsivity while preserving the overall task structure (Henze et al., 2020, 2025). Site-specific study protocols and the timing of cortisol sampling and resting-state fMRI acquisition are shown in Supplementary Figure 1.

Resting-state fMRI was acquired immediately after Scan*STRESS* while participants kept their eyes open and fixated a cross. The acquisition therefore captured a post-stress network state temporally aligned with expected HPA-axis activation rather than an unstressed baseline resting condition. Participants were scanned on Siemens 3T MRI scanners: a Magnetom Prisma in Regensburg and a Trio Tim in Berlin (Siemens Healthcare, Erlangen, Germany). Resting-state acquisition parameters were: TR/TE=2000/30ms, flip angle=90°, FOV=192mm, 37 slices (3mm thickness, 1mm gap), and multiband factor=8 in Regensburg, and TR/TE=2020/25ms, flip angle=80°, FOV=192mm, 37 slices (3mm thickness, 0.60mm gap) in Berlin.

### Cortisol assessment and outcome definition

Salivary cortisol was assessed repeatedly across the experimental session using Salivettes® (Sarstedt, Nümbrecht, Germany), with site-specific sampling schedules shown in Supplementary Figure 1. The pre-stress sample at -1min served as the pre-stress baseline. Following the previous mega analysis (Henze et al., 2025), cortisol increase was defined as the individual peak post-stress cortisol concentration minus the pre-stress baseline sample. The individual post-stress peak was determined from samples collected at +30, +50, and +65min in Regensburg and at +30, +45, and +60min in Berlin. Cortisol increase was selected as the primary endocrine outcome (Dickerson & Kemeny, 2004), as the present study focused on interindividual differences in the magnitude of HPA-axis reactivity to the psychosocial stressor. In contrast, alternative integrated indices such as area under the curve are computed across the full sampling period and are therefore also sensitive to the temporal dynamics of the response, including its latency and recovery (Pruessner et al., 2003).

Cortisol concentrations were quantified using site-specific immunoassay platforms: dissociation-enhanced lanthanide fluorescence immunoassay (DELFIA) in Regensburg and chemiluminescence immunoassay (IBL, Hamburg, Germany) in Berlin. For DELFIA, intra- and inter-assay coefficients of variation ranged from 4.0-6.7% and 7.1-9.0%, respectively; for the chemiluminescence assay, both values were below 6.0%. Because the assay platform was confounded with site and salivary cortisol immunoassays can differ systematically in calibration, antibody specificity, and cross-reactivity with related steroids such as cortisone (Miller et al., 2013), *site* was included as a covariate in inferential and predictive analyses.

### Preprocessing

Resting-state fMRI data were preprocessed using the standardized ENIGMA HALFpipe pipeline (Version 1.2.2; Waller et al., 2022). Preprocessing included brain extraction with SynthSeg (Billot et al., 2023), motion correction, susceptibility distortion correction, coregistration, and spatial normalization via fMRIPrep (Esteban et al., 2019), followed by grand mean scaling, ICA-AROMA-based nuisance correction (Pruim et al., 2015), temporal high-pass filtering at .008Hz, and spatial smoothing with a 4mm full-width at half maximum kernel. Preprocessed images were visually inspected by trained researchers.

For each participant, regional mean time series were extracted from 166 regions defined by the Automated Anatomical Labeling atlas 3v2 (AAL3v2; Rolls et al., 2020). Fisher’s *z*-transformed pairwise correlations were computed between all regional time series, yielding a 166×166 rsFC matrix for each participant. For visualization and interpretation, AAL3v2 regions were additionally mapped onto canonical resting-state networks based on Yeo et al. (2011). The region-to-network correspondence is provided in Supplementary Table 3.

### Data Analysis

Analyses comprised two complementary components. First, NBS (Zalesky et al., 2010) was used to identify connected components of post-stress rsFC associated with acute cortisol increase. Second, multivariate predictive modeling was used to test whether post-stress rsFC predicted cortisol increase in unseen individuals using NBS-Predict (Serin et al., 2021) and, as a complementary benchmark, CPM (Shen et al., 2017).

For NBS, multiple regression models were fitted edge-wise with rsFC connections as dependent variables and cortisol increase as the covariate of interest, controlling for sex/hormonal status, *age*, and *site*. *Sex*/*hormonal status* was modeled using dummy-coded categorical regressors (male as reference), distinguishing males, naturally cycling females tested in the luteal phase, females using hormonal contraceptives, and post-menopausal females. Positive and negative associations were tested separately. A primary component-forming threshold of |*t*|>3.1 was used to define suprathreshold connections, and component-level significance was assessed using 5000 permutations. Component-level *p*-values were considered significant at *p*<.05 and FDR-corrected across positive and negative contrasts. To assess sensitivity to the inherently arbitrary choice of the primary component-forming threshold, analyses were repeated across six *t*-thresholds ranging from liberal to conservative values (*t*=2.0, 2.5, 3.0, 3.1, 3.5, and 4.0). Effect sizes for individual connections within significant components were quantified using 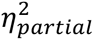 (Cohen, 1973), and nodal strength was calculated from these effect sizes to characterize regional centrality within the identified networks (Supplementary Methods). Additional sex-specific NBS analyses tested male-only, female-only, male>female, and female>male cortisol-rsFC associations; the corresponding general linear model designs are provided in Supplementary Table 4.

For individual-level prediction, we employed NBS-Predict (version v1.0.1; Serin et al., 2021), a framework that integrates NBS-based feature selection with ML within a nested cross-validation (CV) scheme. During each CV iteration, NBS-Predict identifies network components associated with the target variable within the training data and uses these components as predictive features. Selected components (using a threshold of *p*<.01) were then entered into L2-regularized linear regression and support vector regression (SVR) models implemented using *fitrlinear* (MATLAB R2024b). Model performance was evaluated using 10 repetitions of 5-fold outer cross-validation, with 5-fold inner cross-validation for hyperparameter optimization. This procedure enables out-of-sample evaluation and allows the generalizability of cortisol-related connectivity patterns to be assessed, which is not directly provided by conventional NBS (Kriegeskorte et al., 2009). The regularization parameter *λ* was selected within the inner loop from 10 logarithmically spaced values between 10⁻³ and 10⁴. Confounding effects of *age*, *site*, and *sex*/*hormonal status* were removed using cross-validated confound regression (Snoek et al., 2019). Pearson’s correlation coefficient *r* between observed and predicted cortisol increase was used as the primary performance metric. Statistical significance was assessed using 1000 permutations (Ojala & Garriga, 2010), and *p*-values were FDR-corrected across ML algorithms. The primary interpretative output of NBS-Predict was a weighted network indexing the stability and predictive contribution of selected connections across cross-validation folds. For visualization, this network was thresholded at weights of .90 and 1.00 to highlight the most stable and predictive connections. Nodal strength derived from connection weights was used to summarize regional centrality within the predictive network (Supplementary Methods).

Both NBS and NBS-Predict assume that relevant associations are embedded within connected network components, consistent with the view that brain function emerges from interconnected regional networks (Bressler, 1995; Bullmore & Sporns, 2009). CPM (Shen et al., 2017) was applied as a complementary benchmark to assess whether predictive information also resided in connections not constrained to connected components. CPM selected positive and negative predictive edges within the training data, summarized them into network-strength features, and predicted cortisol increase using linear regression. The same cross-validation, confound-regression, performance-evaluation, and permutation-testing framework was used as for NBS-Predict. Full CPM implementation details are provided in the Supplementary Methods.

## RESULTS

### Network-based Statistic

NBS identified a significant component in which post-stress rsFC was negatively associated with acute cortisol increase (258 connections among 78 brain regions, *p_FDR_*=.009, FDR-corrected across positive and negative contrasts; Figure 1 and Supplementary Figure 2). No positively associated component reached significance. The significant negative component was centered on subcortical regions, particularly thalamic nuclei and basal ganglia structures, including the pallidum, putamen, nucleus accumbens (NAcc), ventral tegmental area (VTA), and substantia nigra (SCN). Additional connections extended to DMN regions, including the ACC and PCC, as well as the orbitofrontal cortex, middle cingulate, angular, limbic, and cerebellar regions. Among all regions in the component, the thalamus showed the highest nodal strength, followed by the pallidum and the PCC, indicating that these regions represented central hubs of the cortisol-associated network. The largest individual edge-wise effects were observed for intra-thalamic connections. A complete list of connections and regions is provided in Supplementary Tables 5 and 6.

**Figure 1.**
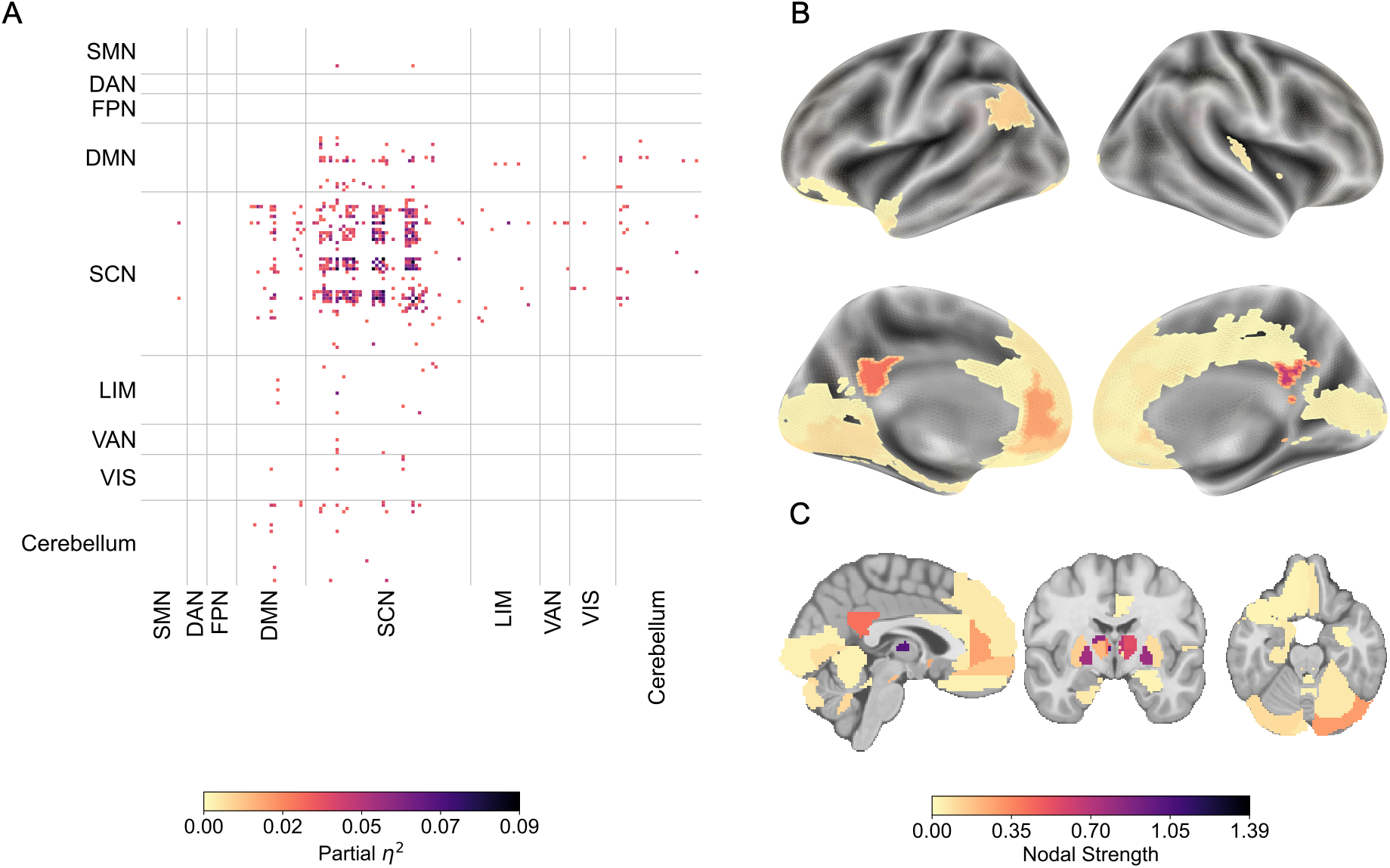
Significant subnetwork (258 connections among 78 regions) negatively associated with cortisol increase revealed by NBS analysis (*t*<-3.1). Positive associations (*t*>3.1) did not reach significance. **A.** The heatmap indicates the intra- and inter-connections, grouped by the assigned canonical functional networks (see Methods for details) and colored according to their effect size (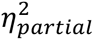). Connections are colored based on their effect size (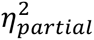). SMN: Somatomotor Network; DAN: Dorsal Attention Network; FPN: Frontoparietal Network; DMN: Default Mode Network; SCN: Subcortical Network; LIM: Limbic Network; VAN: Ventral Attention Network; VIS: Visual Network. **B.** and **C.** Nodal strength of brain regions found in the significant subnetwork is displayed on the brain surface (fsaverage) and in Montréal Neurological Institute (MNI) volumetric space. The details of the nodal strength computation are provided in the Supplementary Methods.

Sensitivity analyses confirmed that the negative cortisol-associated component remained significant across all tested primary *t*-thresholds, demonstrating robustness to the choice of component-forming threshold (Supplementary Table 7, Supplementary Figures 3-7). Across increasingly conservative thresholds, thalamic and pallidal regions, the PCC, left medial orbitofrontal cortex, left parahippocampal gyrus (PHG), and left locus coeruleus (LC) remained consistently represented. In contrast, no positive connected component survived correction at any tested threshold, suggesting that the observed relationship between acute cortisol increase and post-stress rsFC was directionally specific and characterized by lower connectivity in individuals with stronger cortisol response.

Additional sex-stratified analyses revealed a significant negative post-stress rsFC-cortisol association in the female sample after FDR correction (337 connections among 112 brain regions; *p_FDR_*=.011; Figure 2). This component included similar subcortical and PCC hubs as the main analysis, together with more spatially distributed cortical regions that were represented by only one or a few connections. In contrast, the male-only analysis and the formal sex-difference contrasts (male > female, and female>male) did not survive correction (Supplementary Table 8 and Figures 7-9). Thus, although the stratified analyses indicated a significant cortisol-related network in females, there was no robust evidence for a statistically significant difference in cortisol-rsFC associations between males and females.

**Figure 2.**
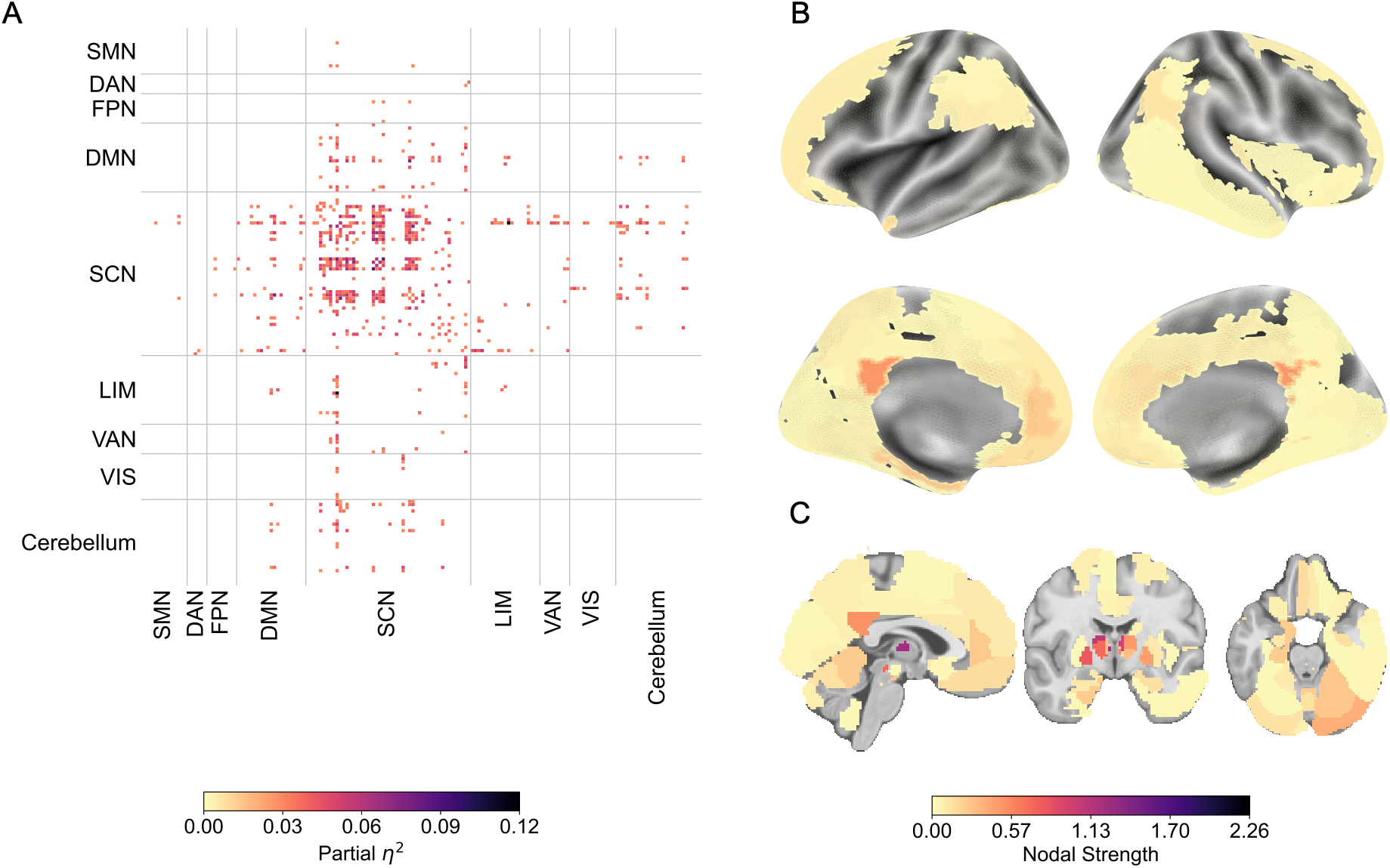
Significant subnetwork (337 connections among 112 regions) negatively associated with cortisol increase in females revealed by NBS analysis (*t*<-3.1). Positive associations (*t*>3.1) and the male sample were not significant after FDR correction. Figures showing the uncorrected results are provided in Supplementary Figures 7-9. **A.** The heatmap indicates the intra- and inter-connections, grouped by the assigned canonical functional networks (see Methods for details) and colored according to their effect size (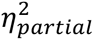). Connections are colored based on their effect size (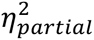). SMN: Somatomotor Network; DAN: Dorsal Attention Network; FPN: Frontoparietal Network; DMN: Default Mode Network; SCN: Subcortical Network; LIM: Limbic Network; VAN: Ventral Attention Network; VIS: Visual Network. **B.** and **C.** Nodal strength of brain regions found in the significant subnetwork is displayed on the brain surface (fsaverage) and in MNI volumetric space. The details of the nodal strength computation were provided in the Supplementary Methods.

### Acute Cortisol Increase Prediction

NBS-Predict yielded modest but significant out-of-sample prediction of acute cortisol increase. Prediction performance was significant for SVR (*r*=.142, *p_uncorr_*=.008, *p_FDR_*=.016) and reached significance for L2-regularized linear regression, although with lower performance (*r*=.117, *p_uncorr_*=.027, *p_FDR_*=.027).

The highest-weight predictive connections, selected across all CV-folds, comprised 27 connections among 15 thalamic subregions and connections between thalamic subregions, the left pallidum, and left medial orbitofrontal cortex (OFC). Additional highly robust connections involved the right PCC, right pallidum, left NAcc, left PHG, and left LC (weight threshold ≥.90; Figure 3), resulting in a robust predictive network of 65 connections among 28 regions. Consistent with the NBS findings, this predictive network was dominated by subcortical connections, particularly involving the thalamus and pallidum, with additional contributions from DMN and limbic regions. All robust predictive connections were negatively associated with acute cortisol increase.

**Figure 3.**
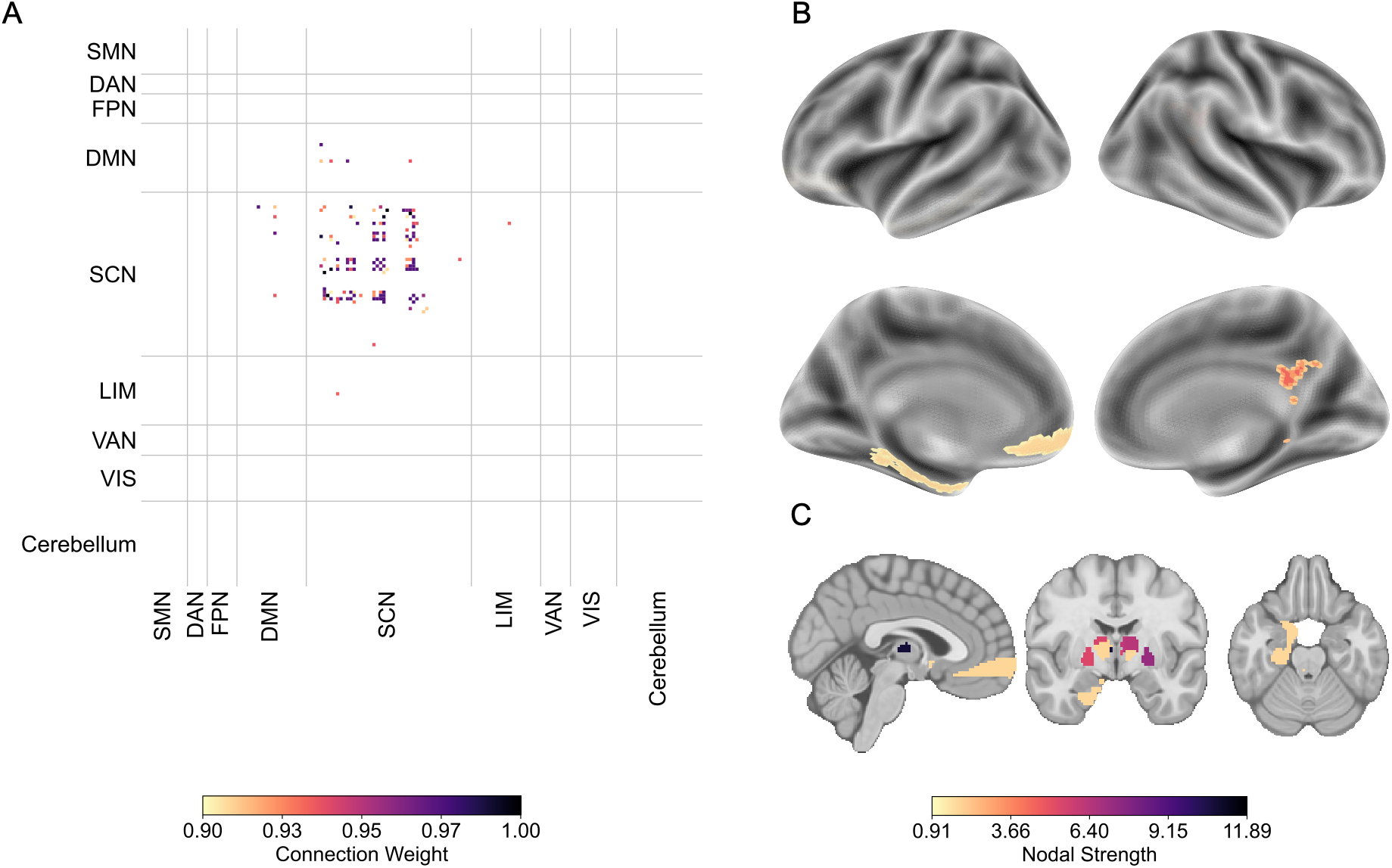
Subnetwork of relevant connections (65 connections among 28 regions), predicting cortisol increase using NBS-Predict (*r*=.142, *p_FDR_*=.016). Weight threshold of .90 was used to visualize the connections with high relevance to the prediction. **A.** The heatmap indicates the intra- and inter-connections within the subnetwork, grouped by the assigned canonical functional networks (see Methods for details) and colored according to their weight, showing their importance. Connections are colored based on their connection weight. SMN: Somatomotor Network; DAN: Dorsal Attention Network; FPN: Frontoparietal Network; DMN: Default Mode Network; SCN: Subcortical Network; LIM: Limbic Network; VAN: Ventral Attention Network; VIS: Visual Network. **B.** and **C.** Nodal strength of brain regions found in the significant subnetwork are displayed on the brain surface (fsaverage) and in MNI volumetric space. The details of the nodal strength computation were provided in the Supplementary Methods.

The complementary CPM analysis also significantly predicted acute cortisol increase. Prediction was significant for the combined network (*r*=.173, *p_FDR_*=.009) and the negative network (*r*=.159, *p_FDR_*=.006), whereas the positive network did not yield significant prediction (*r*=.015, *p_FDR_*=.372). At a selection frequency of ≥0.90, CPM identified a stable set of 78 connections among 33 regions (Supplementary Figure 10). These connections predominantly involved subcortical regions and PCC hubs, showing a pattern of high nodal strength similar to the subnetwork revealed by NBS-Predict. Full lists of connections identified by NBS-Predict and CPM are provided in Supplementary Tables 11-15.

## DISCUSSION

In this study, we investigated how individual differences in acute cortisol increase following psychosocial stress are reflected in post-stress rsFC. Using complementary inferential and predictive approaches, we found that stronger cortisol responses were associated with lower rsFC within a distributed network centered on thalamic nuclei and pallidum and extending to DMN, limbic, ventral attention, and cerebellar regions. The associated network remained significant across multiple NBS thresholds, indicating that the finding was robust to the choice of component-forming threshold. Moreover, NBS-Predict and CPM provided converging evidence that post-stress rsFC contains modest but significant information about individual cortisol reactivity.

A central finding was the consistent negative direction of the association between acute cortisol increase and post-stress rsFC. Participants with stronger cortisol responses showed lower connectivity within the identified network, whereas no positive NBS component survived correction, and predictions across frameworks were driven by the negative rather than the positive network. However, importantly, this pattern should not be interpreted as a global reduction in functional connectivity, but as a specific cortisol-related connectivity phenotype characterized by reduced coupling among stress-relevant subcortical and cortical regions. In this respect, the present findings extend previous stress neuroimaging work (Henze et al., 2023; Henze et al., 2025; Serin et al., 2026) by specifying how endocrine stress reactivity is reflected in the post-stress connectivity state: rather than indicating broad or nonspecific rsFC alteration, acute HPA-axis reactivity was associated with a directionally specific pattern of lower connectivity within a distributed subcortical-cingulate network. By focusing on whole-brain post-stress connectivity and individual-level prediction, this study provides a network-based, predictive perspective on neural-endocrine coupling following acute stress.

The strongest and most consistent involvement of thalamic and pallidal regions suggests that subcortical connectivity is a central feature of the observed cortisol-related post-stress network phenotype. The thalamus is a major hub for cortical-subcortical communication and has previously been implicated in acute stress-related network reconfiguration (Reinelt et al., 2019), while pallidal and adjacent basal ganglia regions are part of circuits involved in motivational, motor, and regulatory processes (Simonyan, 2019; Soares-Cunha & Heinsbroek, 2023). However, given the correlational nature and spatial scale of the present analysis, these functional interpretations should remain cautious. Further, the convergence of thalamic and pallidal involvement across NBS, NBS-Predict, and CPM indicates that cortisol-related post-stress rsFC extends beyond classical limbic stress circuitry and includes broader subcortical network architecture during the post-stress period.

Beyond the thalamic-pallidal core, the cortisol-associated network extended to cortical and cerebellar regions, most notably involving cingulate, orbitofrontal, parahippocampal, and cerebellar connections. The cingulate findings are particularly relevant because PCC and ACC regions are positioned at the interface of default-mode, salience-related, and limbic circuitry and have also been implicated in previous Scan*STRESS* work linking neural stress responses to cortisol reactivity (Henze et al., 2025). Together with recent structural findings from a similar mega-analysis cohort (Henze et al., 2025) associating cingulate morphology with cortisol responses (Serin et al., 2026), the present rsFC results point to cingulate regions as candidate multimodal correlates of HPA-axis stress reactivity. While these cortical regions are previously implicated in stress appraisal, self-referential processing, affective regulation, and recovery (Dimitrov et al., 2018; Kennis et al., 2016; Van Der Werff et al., 2013; Y. Zhang et al., 2020), the interpretation should remain cautious because the present design does not determine whether connectivity of these regions reflects these functions. The cerebellar involvement is also noteworthy but should likewise be viewed as exploratory. Although cerebellar systems are increasingly implicated in affective, cognitive, and autonomic processes (Rudolph et al., 2023; Schmahmann et al., 2019; Snow et al., 2014), their role in psychosocial stress reactivity remains less established. Their involvement here suggests that cortisol-related post-stress connectivity may extend beyond classical limbic, subcortical, and cortical circuits, but future studies are needed to determine whether cerebellar connectivity is a reliable component of neural-endocrine stress coupling.

Given previous evidence that neural and endocrine stress responses may vary by sex (Henze et al., 2021, 2025; Kogler et al., 2016), we additionally examined this assumption in sex-stratified NBS analyses. Here, a significant negative cortisol-rsFC association was observed in the female sample (*n*=193), involving a network that overlapped with the full-sample findings in thalamic, subcortical, and PCC regions, while also extending across additional cortical areas. However, neither the male subgroup (*n*=146) nor the formal sex-difference contrasts survived correction. Notably, this is unlikely to be attenuated or restricted endocrine reactivity: cortisol increases were, in fact, larger in males (5.10±5.28 vs. 2.60±5.38nmol/L, *d*=0.47), with near-identical dispersion in both subgroups. In addition, the male subgroup was also only modestly smaller than the female subgroup (146 vs. 193 participants). This highlights the need for future studies that are specifically powered to test whether these patterns reflect robust sex-specific effects, hormonal-status effects, or sample-specific variability. Furthermore, whereas previous structural studies using similar cohorts predominantly identified brain-stress associations in males (Henze et al., 2023; Serin et al., 2026), the present female subgroup finding suggests that sex-related variability in neural-endocrine coupling may differ across imaging modalities. Multimodal neuroimaging approaches may therefore be particularly informative for disentangling whether sex-related effects are modality-specific, reflect distinct neural mechanisms, or arise from differences in sample composition and hormonal-status distributions.

NBS-Predict achieved a modest (*r*=.142) but significant prediction of acute cortisol increase from post-stress rsFC, and CPM yielded slightly higher prediction performance (*r*=.173). This difference may reflect the methodological distinction between NBS-Predict, which prioritizes connected network components, and CPM, which can retain predictive information in individual edges irrespective of their topological embedding. Although direct comparison with previous ML work is not straightforward, W. Zhang et al. (2020) similarly suggested that stress-related rsFC contains information about cortisol responses, albeit by relating cortisol increase to classification probabilities distinguishing stress from rest rather than by directly predicting cortisol increase. Although predictive effect sizes were small, the overlap between the inferential NBS component and the predictive networks supports the internal consistency of the observed cortisol-related connectivity pattern. A consistent core network, comprising the PCC, medial OFC, thalamus, and pallidum, was implicated across predictive frameworks, with CPM additionally capturing isolated cerebellar and subcortical connections, which suggest that these regions contributed consistently to cortisol-related post-stress rsFC within the present dataset. However, because these analyses were conducted in the same sample, this convergence should be interpreted as within-sample methodological consistency rather than independent replication. Thus, the external consistency (or robustness) of the results should be evaluated across multiple datasets in future studies.

Several limitations should be considered. First, although the study focused on post-stress rsFC because of its temporal alignment with HPA-axis activation, post-stress rsFC is not independent of the brain’s intrinsic functional architecture. Instead, it likely reflects a combination of stable interindividual differences and acute stress-reactive reorganization. Therefore, the separation of stress-induced changes in connectivity from pre-existing interindividual differences remains limited. Although such separation would require the acquisition of both pre- and post-stress rsFC and may still be affected by the limited reliability of resting-state fMRI (Noble et al., 2019), it remains an important direction for future studies. Second, as the present study focused specifically on cortisol increase, the findings should be interpreted as reflecting a cortisol-related post-stress connectivity phenotype rather than a general neural signature of stress reactivity. Other stress-response dimensions, including autonomic, affective, and behavioral responses, as well as neuroendocrine measures related to HPA-axis reactivity, may show partially overlapping yet distinct neural correlates. Third, rs-fMRI acquisition length differed across sites, with a substantially shorter acquisition in Berlin (5.18min) than in Regensburg (18min). Acquisition length may affect not only the precision and reliability of functional connectivity estimates, but also the extent to which the scan captures the evolving post-stress HPA-axis response. Although we accounted for site-related variation in the current analyses, residual site- or acquisition-related differences, as well as what to capture, cannot be fully excluded and should be considered when interpreting the results. Nevertheless, the supplementary re-analysis with truncated rs-fMRI data from the Regensburg site, which matches the acquisition length in Berlin, demonstrated similar results (Supplementary Figure 11). Fourth, despite statistically significant prediction, predictive performance was modest. The present findings therefore do not support strong claims regarding clinically useful predictive biomarkers of cortisol stress reactivity. Instead, they suggest that post-stress rsFC captures a statistically reliable but limited proportion of interindividual variability in acute cortisol increase. This is consistent with the broader view that endocrine stress responses are shaped by multiple interacting neural, physiological, psychological, and contextual factors (Herman et al., 2016; Kudielka et al., 2009).

In conclusion, the present data-driven findings suggest that acute endocrine stress response after psychosocial stress exposure is associated with a distributed post-stress connectivity phenotype characterized mainly by lower coupling among subcortical, DMN, limbic, and cerebellar regions. The convergence of NBS, NBS-Predict, and CPM indicates that post-stress rsFC contains consistent inferential and predictive information about individual HPA-axis reactivity, although predictive effects were modest. These findings support a network-level view of neural-endocrine coupling after acute stress and highlight subcortical-cingulate connectivity as a promising target for future work on stress susceptibility and resilience.

## ACKNOWLEDGEMENTS

We thank Franziska Bauer, Patricia Bohmann, Zora-Pia Gerbers, Tanja Julia Hiltl, Susanne Kargl, Elias Luigi Landfried, Daniel Schleicher, Katharina Spannruft, Philipp Vogel, Marina Weber, Michael Zeitler, and Florian Zimmer for their help with running the experimental sessions and data entry in Regensburg. Additionally, we thank all those who assisted with the initial data collection for the original Berlin site study.

## FUNDING RESOURCES

This study was supported by German Research Foundation Grants No. HE9212/1-1 (to G-IH, project number 513531314). Further support for the original data collection in Regensburg was provided by grant numbers: KU1401/6-1, KU1401/9-1, KU1401/9-2, KU1401/10-1, WU392/9-1 [to BMK and SW].

## DATA AVAILABILITY STATEMENT

The datasets analyzed during the current study are available from the corresponding author upon reasonable request. The code used to analyze the data and generate the presented figures can be found at: https://github.com/eminSerin/stress-rs-paper. A repository of studies that have already used and published the data from the Regensburg site is available at: https://osf.io/echja/.

## AUTHOR CONTRIBUTIONS

**Emin Serin**: Conceptualization, Methodology, Software, Validation, Formal Analysis, Data Curation, Writing – Original Draft, Writing – Review & Editing, Visualization. **Elvan Emurla**: Data Curation, Writing – Original Draft. **Christoph Bärtl**: Investigation, Writing – Review & Editing. **Marina Giglberger**: Investigation, Writing – Review & Editing. **Julian Konzok**: Investigation, Writing – Review & Editing. **Hannah L. Peter**: Investigation. **Ludwig Kreuzpointner**: Writing – Review & Editing. **Brigitte M. Kudielka**: Funding Acquisition, Writing – Review & Editing. **Stefan Wüst**: Funding Acquisition, Writing – Review & Editing. **Susanne Erk**: Data Curation, Resources, Writing – Review & Editing. **Henrik Walter**: Resources, Writing – Review & Editing, Supervision, Funding Acquisition. **Gina-Isabelle Henze**: Conceptualization, Methodology, Software, Validation, Formal Analysis, Investigation, Resources, Data Curation, Writing – Original Draft, Writing – Review & Editing, Visualization, Project Administration, Supervision, Funding Acquisition.

## DECLARATION OF GENERATIVE AI USE

During the preparation of this work, the authors used ChatGPT (OpenAI) and Claude (Anthropic) to support language editing, structural refinement, and drafting assistance, as well as to develop analysis scripts. After using these tools, the authors reviewed and edited the content as needed and took full responsibility for the content of the published article.

