## Supplementary Material for "Linking post-stress brain connectivity to acute cortisol reactivity using network-based inference and prediction"

### **Supplementary Methods**

**Sample Inclusion Criteria**

Across studies, inclusion criteria required the absence of current psychiatric, neurological, or endocrine disorders and no use of medication affecting the central nervous system or endocrine function. Participants were additionally required to smoke fewer than five cigarettes per day, abstain from daily alcohol consumption, meet magnetic resonance imaging (MRI) safety criteria (e.g., no metal implants, not pregnant), report no current major stressful life event, and not engage in night shift work.

**Nodal Degree and Nodal Strength**

Nodal degree was defined as the number of connections (i.e., edges) incident on a given region (i.e., node), with higher values indicating greater participation in network connections and potential hub status (Bullmore & Sporns, 2009; Fornito et al., 2016). Nodal strength, the weighted analog of nodal degree, was defined as the sum of the weights of all connections incident on a region (Bullmore & Sporns, 2009; Fornito et al., 2016). The connection weight used to compute nodal strength in the NBS analysis was $\eta_{partial}^{2}$ the effect size, while the weights within the output network and the selection‑frequency values served as connection weights in the NBS‑Predict and CPM analyses.

**Connectome-based Predictive Modeling (CPM)**

As a complementary analysis to NBS-Predict, CPM (Shen et al., 2017) was applied to examine whether dispersed connections were associated with acute cortisol increase and to benchmark prediction performance against the component-based approach. Within each fold of a 10-repetition, 5-fold cross-validation, confounding effects of age, site, and the combined sex-and-hormonal-status variable were first removed using cross-validated confound regression (Snoek et al., 2019). Resting-state functional connections were then correlated with acute cortisol increase using Pearson's correlation coefficient, and those surviving a threshold of *p*<.01 were retained. Connections negatively or positively correlated with acute cortisol increase were assigned to negative or positive networks, respectively; these were further merged into a combined network. The summed FC strength of connections within each network (negative, positive, and combined) served as a summary score to predict acute cortisol increase via linear regression. Pearson's correlation coefficient was used as the primary performance metric across all three models, and significance was evaluated via permutation testing (1000 permutations; Ojala & Garriga, 2010) and FDR correction. The contribution of individual connections to the predictive model was assessed by their stability (i.e., selection frequency) across CV folds.

### **Supplementary Tables**

| **Supplementary Table 1.** Demographics and cortisol increase across acquisition sites. | | | |
| --- | --- | --- | --- |
| **Variable (µ ± σ)** | **Site** | | ***p*-value** |
|  | **Regensburg** | **Berlin** |  |
| Sex male/female | 113/148 | 33/45 | .98^a^ |
| Age (in years) | 26.24±9.01 | 28.04±6.65 | .04^b^ |
| Cortisol (nmol/L) | 2.80±3.72 | 6.61±8.57 | .00 ^b^ |
| **Note:** ^a^Chi-square test; ^b^Permutation-tested t-test | | | |

| **Supplementary Table 2.** Cortisol across sex and hormonal status. | | | |
| --- | --- | --- | --- |
| **Variable (µ ± σ)** |  | **Cortisol Increase**  **(µ±σ)** | ***p*-value** |
| Sex | Male (*n*=146) | 5.10±5.28 | .00^a^ |
|  | Female (*n*=193) | 2.60±5.38 |  |
| Hormonal Status | Luteal (*n*=102) | 2.97±6.32 | .80^b^ |
|  | Contraceptive (*n*=80) | 2.23±4.26 |  |
|  | Post-menopausal (*n*=11) | 1.90±2.66 |  |
| **Note:** ^a^Permutation-tested t-test; ^b^Permutation-tested one-way ANOVA | | | |

| **Supplementary Table 3.** List of brain regions from AAL3 atlas (Rolls et al., 2020) and the corresponding large-scale Yeo (Yeo et al., 2011) and Triple networks (Van Oort et al., 2017). | | | | | |
| --- | --- | --- | --- | --- | --- |
| **AAL Label** | **Yeo Network** | **Triple Network** | **AAL Label** | **Yeo Network** | **Triple Network** |
| Precentral_L | SMN | - | Temporal_Pole_Sup_R | LIM | SN |
| Precentral_R | SMN | - | Temporal_Mid_L | DMN | SN |
| Frontal_Sup_2_L | DMN | CEN | Temporal_Mid_R | DMN | SN |
| Frontal_Sup_2_R | DMN | CEN | Temporal_Pole_Mid_L | LIM | SN |
| Frontal_Mid_2_L | FPN | CEN | Temporal_Pole_Mid_R | LIM | SN |
| Frontal_Mid_2_R | FPN | CEN | Temporal_Inf_L | LIM | SN |
| Frontal_Inf_Oper_L | VAN | CEN | Temporal_Inf_R | LIM | SN |
| Frontal_Inf_Oper_R | FPN | CEN | Cerebellum_Crus1_L | Cerebellum | - |
| Frontal_Inf_Tri_L | FPN | SN | Cerebellum_Crus1_R | Cerebellum | - |
| Frontal_Inf_Tri_R | FPN | SN | Cerebellum_Crus2_L | Cerebellum | - |
| Frontal_Inf_Orb_2_L | DMN | SN | Cerebellum_Crus2_R | Cerebellum | - |
| Frontal_Inf_Orb_2_R | DMN | SN | Cerebellum_3_L | Cerebellum | - |
| Rolandic_Oper_L | SMN | - | Cerebellum_3_R | Cerebellum | - |
| Rolandic_Oper_R | SMN | - | Cerebellum_4_5_L | Cerebellum | - |
| Supp_Motor_Area_L | SMN | - | Cerebellum_4_5_R | Cerebellum | - |
| Supp_Motor_Area_R | SMN | - | Cerebellum_6_L | Cerebellum | - |
| Olfactory_L | LIM | - | Cerebellum_6_R | Cerebellum | - |
| Olfactory_R | LIM | - | Cerebellum_7b_L | Cerebellum | - |
| Frontal_Sup_Medial_L | DMN | DMN | Cerebellum_7b_R | Cerebellum | - |
| Frontal_Sup_Medial_R | DMN | DMN | Cerebellum_8_L | Cerebellum | - |
| Frontal_Med_Orb_L | DMN | DMN | Cerebellum_8_R | Cerebellum | - |
| Frontal_Med_Orb_R | DMN | DMN | Cerebellum_9_L | Cerebellum | - |
| Rectus_L | LIM | DMN | Cerebellum_9_R | Cerebellum | - |
| Rectus_R | LIM | DMN | Cerebellum_10_L | Cerebellum | - |
| OFCmed_L | LIM | DMN | Cerebellum_10_R | Cerebellum | - |
| OFCmed_R | LIM | DMN | Vermis_1_2 | Cerebellum | - |
| OFCant_L | LIM | DMN | Vermis_3 | Cerebellum | - |
| OFCant_R | FPN | DMN | Vermis_4_5 | Cerebellum | - |
| OFCpost_L | LIM | DMN | Vermis_6 | Cerebellum | - |
| OFCpost_R | LIM | DMN | Vermis_7 | Cerebellum | - |
| OFClat_L | DMN | DMN | Vermis_8 | Cerebellum | - |
| OFClat_R | DMN | DMN | Vermis_9 | Cerebellum | - |
| Insula_L | VAN | SN | Vermis_10 | Cerebellum | - |
| Insula_R | VAN | SN | Thal_AV_L | SCN | SN |
| Cingulate_Mid_L | VAN | CEN | Thal_AV_R | SCN | SN |
| Cingulate_Mid_R | VAN | CEN | Thal_LP_L | SCN | SN |
| Cingulate_Post_L | DMN | DMN | Thal_LP_R | SCN | SN |
| Cingulate_Post_R | DMN | DMN | Thal_VA_L | SCN | SN |
| Hippocampus_L | LIM | DMN | Thal_VA_R | SCN | SN |
| Hippocampus_R | LIM | DMN | Thal_VL_L | SCN | SN |
| ParaHippocampal_L | LIM | DMN | Thal_VL_R | SCN | SN |
| ParaHippocampal_R | LIM | DMN | Thal_VPL_L | SCN | SN |
| Amygdala_L | LIM | SN | Thal_VPL_R | SCN | SN |
| Amygdala_R | LIM | SN | Thal_IL_L | SCN | SN |
| Calcarine_L | VIS | - | Thal_IL_R | SCN | SN |
| Calcarine_R | VIS | - | Thal_Re_L | SCN | SN |
| Cuneus_L | VIS | - | Thal_Re_R | SCN | SN |
| Cuneus_R | VIS | - | Thal_MDm_L | SCN | SN |
| Lingual_L | VIS | - | Thal_MDm_R | SCN | SN |
| Lingual_R | VIS | - | Thal_MDl_L | SCN | SN |
| Occipital_Sup_L | VIS | - | Thal_MDl_R | SCN | SN |
| Occipital_Sup_R | VIS | - | Thal_LGN_L | SCN | SN |
| Occipital_Mid_L | VIS | - | Thal_LGN_R | SCN | SN |
| Occipital_Mid_R | VIS | - | Thal_MGN_L | SCN | SN |
| Occipital_Inf_L | VIS | - | Thal_MGN_R | SCN | SN |
| Occipital_Inf_R | VIS | - | Thal_PuI_L | SCN | SN |
| Fusiform_L | VIS | - | Thal_PuI_R | SCN | SN |
| Fusiform_R | VIS | - | Thal_PuM_L | SCN | SN |
| Postcentral_L | SMN | - | Thal_PuM_R | SCN | SN |
| Postcentral_R | SMN | - | Thal_PuA_L | SCN | SN |
| Parietal_Sup_L | DAN | CEN | Thal_PuA_R | SCN | SN |
| Parietal_Sup_R | DAN | CEN | Thal_PuL_L | SCN | SN |
| Parietal_Inf_L | DAN | DMN | Thal_PuL_R | SCN | SN |
| Parietal_Inf_R | FPN | DMN | ACC_sub_L | DMN | DMN |
| SupraMarginal_L | VAN | SN | ACC_sub_R | DMN | DMN |
| SupraMarginal_R | VAN | SN | ACC_pre_L | DMN | DMN |
| Angular_L | DMN | DMN/CEN | ACC_pre_R | DMN | DMN |
| Angular_R | DMN | DMN/CEN | ACC_sup_L | VAN | SN |
| Precuneus_L | DMN | DMN | ACC_sup_R | VAN | SN |
| Precuneus_R | DAN | DMN | N_Acc_L | SCN | SN/LIM |
| Paracentral_Lobule_L | SMN | - | N_Acc_R | SCN | SN/LIM |
| Paracentral_Lobule_R | SMN | - | VTA_L | SCN | - |
| Caudate_L | SCN | SN/LIM | VTA_R | SCN | - |
| Caudate_R | SCN | SN/LIM | SN_pc_L | SCN | - |
| Putamen_L | SCN | SN/LIM | SN_pc_R | SCN | - |
| Putamen_R | SCN | SN/LIM | SN_pr_L | SCN | - |
| Pallidum_L | SCN | SN/LIM | SN_pr_R | SCN | - |
| Pallidum_R | SCN | SN/LIM | Red_N_L | SCN | - |
| Heschl_L | SMN | - | Red_N_R | SCN | - |
| Heschl_R | SMN | - | LC_L | SCN | - |
| Temporal_Sup_L | SMN | SN | LC_R | SCN | - |
| Temporal_Sup_R | SMN | SN | Raphe_D | SCN | - |
| Temporal_Pole_Sup_L | LIM | SN | Raphe_M | SCN | - |
| **Note:** FC: Functional connectivity. SMN: Somatomotor Network; FPN: Frontoparietal Network; VAN: Ventral Attention Network; DMN: Default Mode Network; DAN: Dorsal Attention Network; LIM: Limbic Network; VIS: Visual Network; SCN: Subcortical Network; SN: Salience Network; CEN: Central Executive Network | | | | | |

| **Supplementary Table 4.** Sex-specific models tested by NBS. | | | | | |
| --- | --- | --- | --- | --- | --- |
| **Contrast** | **Male Cortisol** | **Female Cortisol** | **Site** | **Cycle** | **Age** |
| Male Negative | -1 | 0 | 0 | 0 | 0 |
| Female Negative | 0 | -1 | 0 | 0 | 0 |
| Male Positive | 1 | 0 | 0 | 0 | 0 |
| Female Positive | 0 | 1 | 0 | 0 | 0 |
| Male > Female | 1 | -1 | 0 | 0 | 0 |
| Female > Male | -1 | 1 | 0 | 0 | 0 |
| **Note:** Negative and positive models reveal negative and positive rsFC‑cortisol associations. | | | | | |

| **Supplementary Table 5.** Connections within a network negatively associated with acute cortisol increase, identified by NBS. | | | |
| --- | --- | --- | --- |
| **Region A** | **Region B** | ***t*** | $\boldsymbol{\eta}_{\boldsymbol{partial}}^{\boldsymbol{2}}$ |
| Thal_MDm_L | Thal_MDl_R | 5.743 | .090 |
| Thal_MDm_R | Thal_PuA_L | 5.529 | .084 |
| Thal_VPL_L | Thal_MDm_L | 5.371 | .080 |
| Thal_LP_R | Thal_MDl_R | 5.363 | .080 |
| Thal_PuA_L | Thal_PuA_R | 5.329 | .079 |
| Thal_MDl_R | Thal_PuM_L | 5.289 | .078 |
| Thal_LP_R | Thal_MDm_L | 5.149 | .074 |
| Thal_VA_L | Thal_PuA_L | 5.118 | .073 |
| Thal_VL_R | Thal_PuA_L | 5.089 | .072 |
| Thal_VPL_L | Thal_MDl_R | 5.059 | .072 |
| Thal_VL_R | Thal_PuM_L | 4.900 | .067 |
| Thal_MDl_R | Thal_PuA_L | 4.894 | .067 |
| Thal_MDl_R | Thal_PuM_R | 4.885 | .067 |
| Thal_MDm_L | Thal_PuA_L | 4.874 | .067 |
| Thal_MDm_L | Thal_MDm_R | 4.870 | .067 |
| Thal_MDl_L | Thal_PuA_L | 4.838 | .066 |
| Thal_VL_L | Thal_MDm_L | 4.823 | .065 |
| Thal_VL_L | Thal_MDl_R | 4.823 | .065 |
| Thal_MDl_R | Thal_PuA_R | 4.820 | .065 |
| Thal_VPL_L | Thal_MDm_R | 4.724 | .063 |
| Thal_MDm_R | Thal_MDl_L | 4.723 | .063 |
| Thal_MDl_L | Thal_MDl_R | 4.646 | .061 |
| Pallidum_R | Thal_PuI_R | 4.575 | .059 |
| Thal_PuM_R | Thal_PuA_L | 4.549 | .059 |
| Thal_VA_R | Thal_PuM_L | 4.543 | .059 |
| Thal_VPL_L | Thal_PuA_R | 4.539 | .058 |
| Cingulate_Post_R | Thal_PuM_R | 4.531 | .058 |
| Thal_AV_L | Thal_PuM_R | 4.527 | .058 |
| ParaHippocampal_L | Thal_LP_R | 4.526 | .058 |
| Thal_AV_R | Thal_MDm_L | 4.523 | .058 |
| Thal_AV_R | Thal_PuA_L | 4.518 | .058 |
| Thal_VL_L | Thal_PuA_L | 4.507 | .058 |
| Thal_VA_R | Thal_MDm_L | 4.478 | .057 |
| Cingulate_Post_R | Thal_VL_L | 4.466 | .057 |
| Thal_VPL_L | Thal_PuM_R | 4.464 | .057 |
| Thal_LP_R | Thal_PuA_R | 4.458 | .056 |
| Thal_AV_R | Thal_VPL_L | 4.456 | .056 |
| Thal_VL_L | Thal_MDm_R | 4.445 | .056 |
| Thal_AV_R | Thal_MDl_R | 4.442 | .056 |
| Pallidum_R | Thal_PuA_L | 4.441 | .056 |
| Thal_VL_R | Thal_MDm_L | 4.411 | .055 |
| Pallidum_R | Thal_LGN_L | 4.384 | .055 |
| Thal_AV_R | Thal_PuM_L | 4.377 | .055 |
| Thal_MDl_L | Thal_PuM_L | 4.375 | .055 |
| Frontal_Med_Orb_L | Pallidum_L | 4.374 | .054 |
| Thal_LP_R | Thal_PuA_L | 4.361 | .054 |
| Thal_PuA_R | Thal_PuL_R | 4.325 | .053 |
| Cingulate_Post_R | Thal_AV_R | 4.323 | .053 |
| Thal_MDm_L | Thal_PuM_L | 4.307 | .053 |
| Thal_MDl_L | Thal_PuM_R | 4.283 | .052 |
| Pallidum_R | Thal_PuM_R | 4.273 | .052 |
| Thal_LP_R | Thal_VPL_R | 4.265 | .052 |
| Thal_LP_R | Thal_MDl_L | 4.223 | .051 |
| Thal_MDm_L | Thal_PuM_R | 4.213 | .051 |
| Thal_AV_R | Thal_VL_R | 4.209 | .051 |
| Thal_IL_L | Thal_PuM_R | 4.192 | .050 |
| Thal_VL_R | Thal_MDl_R | 4.175 | .050 |
| Thal_PuM_R | Thal_PuL_R | 4.156 | .049 |
| Thal_MDm_L | LC_L | 4.147 | .049 |
| Pallidum_L | Thal_VL_R | 4.110 | .048 |
| Thal_VA_L | Thal_VA_R | 4.102 | .048 |
| Thal_VPL_L | Thal_PuM_L | 4.094 | .048 |
| Pallidum_L | Pallidum_R | 4.073 | .048 |
| Thal_MDm_L | Thal_PuL_L | 4.068 | .047 |
| Pallidum_R | Thal_PuM_L | 4.063 | .047 |
| Cingulate_Post_L | Thal_PuA_L | 4.040 | .047 |
| Thal_AV_R | Thal_VA_L | 4.038 | .047 |
| Thal_MDm_L | Thal_PuA_R | 3.997 | .046 |
| Thal_MDl_R | Thal_LGN_L | 3.985 | .046 |
| Pallidum_L | Thal_MDm_L | 3.984 | .046 |
| Thal_PuM_L | Thal_PuA_L | 3.976 | .045 |
| Thal_VA_R | Thal_PuM_R | 3.946 | .045 |
| Thal_AV_R | Thal_MDm_R | 3.942 | .045 |
| Temporal_Pole_Mid_L | Thal_PuL_L | 3.929 | .044 |
| Thal_VL_L | Thal_PuM_R | 3.928 | .044 |
| Thal_PuL_R | N_Acc_L | 3.925 | .044 |
| Pallidum_R | Thal_MDm_L | 3.920 | .044 |
| Cingulate_Post_L | VTA_L | 3.910 | .044 |
| Thal_LP_R | ACC_pre_L | 3.901 | .044 |
| Thal_AV_L | Thal_PuA_L | 3.901 | .044 |
| Thal_LP_R | Thal_MDm_R | 3.885 | .043 |
| Putamen_L | Thal_LP_R | 3.877 | .043 |
| Vermis_9 | Thal_LGN_L | 3.862 | .043 |
| Cerebellum_Crus2_R | Thal_PuA_L | 3.831 | .042 |
| Cerebellum_Crus1_R | Thal_PuA_L | 3.807 | .042 |
| Cingulate_Post_R | Pallidum_L | 3.796 | .042 |
| Thal_VPL_L | Thal_PuA_L | 3.790 | .041 |
| Cingulate_Post_R | Thal_LGN_L | 3.777 | .041 |
| Thal_IL_R | ACC_sub_R | 3.770 | .041 |
| Thal_VL_L | Thal_MDl_L | 3.753 | .041 |
| Pallidum_L | Thal_MDl_L | 3.739 | .040 |
| Cingulate_Post_L | Cerebellum_Crus1_R | 3.724 | .040 |
| Thal_VA_R | ACC_sub_R | 3.722 | .040 |
| Thal_VPL_L | Thal_MGN_L | 3.699 | .040 |
| Thal_VPL_R | Thal_MDl_R | 3.698 | .040 |
| Thal_VPL_L | Thal_MDl_L | 3.698 | .040 |
| Heschl_R | Thal_LP_R | 3.695 | .040 |
| Thal_LP_R | N_Acc_L | 3.694 | .039 |
| Thal_PuM_L | Thal_PuL_R | 3.693 | .039 |
| Thal_PuI_R | Thal_PuA_L | 3.689 | .039 |
| Thal_VL_L | Thal_VL_R | 3.689 | .039 |
| Thal_MDl_L | Thal_PuA_R | 3.685 | .039 |
| Thal_MDm_R | Thal_PuM_R | 3.684 | .039 |
| Calcarine_R | Thal_PuI_R | 3.678 | .039 |
| Thal_LP_R | Thal_PuM_L | 3.675 | .039 |
| Cerebellum_Crus1_R | Thal_MDl_R | 3.673 | .039 |
| Cingulate_Post_R | VTA_L | 3.665 | .039 |
| Thal_Re_R | ACC_pre_L | 3.661 | .039 |
| Thal_IL_L | Thal_PuM_L | 3.654 | .039 |
| Thal_PuA_L | Thal_PuL_L | 3.653 | .039 |
| Pallidum_L | Thal_VPL_R | 3.651 | .039 |
| Thal_AV_R | Thal_PuA_R | 3.642 | .038 |
| Thal_VL_L | Thal_PuM_L | 3.640 | .038 |
| Frontal_Med_Orb_L | Thal_LP_R | 3.630 | .038 |
| Vermis_1_2 | Thal_Re_L | 3.628 | .038 |
| Cingulate_Post_R | Thal_VA_R | 3.624 | .038 |
| Thal_VL_R | Thal_PuA_R | 3.616 | .038 |
| Frontal_Med_Orb_R | Pallidum_L | 3.614 | .038 |
| Thal_PuA_L | ACC_pre_L | 3.598 | .038 |
| Cingulate_Post_R | Thal_PuA_L | 3.596 | .037 |
| Pallidum_L | N_Acc_L | 3.579 | .037 |
| Thal_LP_R | Thal_IL_R | 3.577 | .037 |
| Thal_AV_R | Thal_VL_L | 3.577 | .037 |
| Thal_AV_L | Thal_PuM_L | 3.572 | .037 |
| Thal_VA_L | Thal_PuM_L | 3.572 | .037 |
| Thal_VPL_R | Thal_PuM_R | 3.571 | .037 |
| Thal_VPL_R | Thal_MDm_R | 3.554 | .037 |
| Thal_VPL_R | Thal_PuM_L | 3.552 | .037 |
| Thal_PuM_L | N_Acc_L | 3.548 | .037 |
| Putamen_L | Thal_PuM_R | 3.545 | .036 |
| Thal_MDm_R | Thal_PuM_L | 3.537 | .036 |
| Putamen_R | Thal_LP_R | 3.534 | .036 |
| Pallidum_R | Thal_PuI_L | 3.532 | .036 |
| Cingulate_Post_L | Thal_AV_R | 3.523 | .036 |
| Cingulate_Post_R | Cerebellum_Crus1_R | 3.523 | .036 |
| Cerebellum_Crus1_L | Thal_PuA_L | 3.522 | .036 |
| Thal_MDl_R | Thal_PuL_L | 3.521 | .036 |
| Frontal_Med_Orb_L | VTA_L | 3.518 | .036 |
| Angular_L | Thal_LGN_L | 3.501 | .036 |
| Thal_LP_R | LC_R | 3.498 | .036 |
| Thal_VPL_R | Thal_PuA_R | 3.497 | .036 |
| Thal_VA_R | Thal_VPL_L | 3.491 | .035 |
| Rectus_L | VTA_L | 3.491 | .035 |
| Pallidum_R | Thal_VL_L | 3.487 | .035 |
| Pallidum_L | Cerebellum_Crus2_R | 3.478 | .035 |
| Cingulate_Post_R | Vermis_4_5 | 3.476 | .035 |
| Thal_LP_R | ACC_sup_L | 3.459 | .035 |
| Thal_VA_R | Thal_MDl_L | 3.456 | .035 |
| Cingulate_Post_R | Thal_VPL_L | 3.452 | .035 |
| Thal_MDm_R | Thal_MDl_R | 3.446 | .035 |
| Thal_PuM_R | SN_pc_L | 3.445 | .035 |
| Cingulate_Post_L | Cerebellum_4_5_R | 3.428 | .034 |
| Pallidum_R | Thal_MDl_R | 3.427 | .034 |
| Cerebellum_Crus1_L | Thal_MDl_R | 3.426 | .034 |
| Vermis_9 | Thal_AV_R | 3.426 | .034 |
| Cingulate_Post_R | Thal_MDl_L | 3.421 | .034 |
| Thal_VA_R | Thal_PuA_L | 3.419 | .034 |
| Calcarine_L | Thal_PuI_R | 3.411 | .034 |
| Thal_AV_L | Thal_MDm_R | 3.407 | .034 |
| Cerebellum_Crus1_R | Thal_PuL_L | 3.393 | .034 |
| Cingulate_Post_R | Thal_MDm_L | 3.393 | .034 |
| Cerebellum_Crus1_R | Thal_LP_R | 3.389 | .033 |
| Thal_PuI_R | Thal_PuM_R | 3.377 | .033 |
| Cingulate_Post_L | Cerebellum_6_R | 3.376 | .033 |
| Thal_VL_R | Thal_VPL_L | 3.373 | .033 |
| Pallidum_R | Thal_LGN_R | 3.367 | .033 |
| Rectus_R | VTA_R | 3.365 | .033 |
| Pallidum_R | Thal_PuA_R | 3.362 | .033 |
| Thal_AV_R | Thal_PuM_R | 3.361 | .033 |
| Thal_VL_L | Thal_VPL_L | 3.360 | .033 |
| Pallidum_R | Cerebellum_Crus1_R | 3.360 | .033 |
| Cingulate_Post_L | Thal_VL_L | 3.332 | .032 |
| Thal_AV_L | Thal_MDl_R | 3.331 | .032 |
| Pallidum_R | Thal_MDl_L | 3.322 | .032 |
| Thal_MDl_R | ACC_sup_R | 3.315 | .032 |
| Thal_VL_L | Thal_PuA_R | 3.312 | .032 |
| Amygdala_R | Angular_L | 3.303 | .032 |
| Cerebellum_6_R | Thal_LP_R | 3.303 | .032 |
| Thal_VA_R | Thal_VL_R | 3.302 | .032 |
| Lingual_L | Thal_PuI_R | 3.301 | .032 |
| Pallidum_L | Thal_VPL_L | 3.300 | .032 |
| Putamen_R | Thal_PuM_L | 3.297 | .032 |
| Pallidum_L | Thal_VL_L | 3.295 | .032 |
| Pallidum_R | Thal_VPL_L | 3.293 | .032 |
| Thal_VA_R | ACC_pre_R | 3.285 | .031 |
| Pallidum_L | ACC_pre_L | 3.285 | .031 |
| Thal_MDl_R | Thal_MGN_L | 3.284 | .031 |
| Pallidum_L | Thal_AV_R | 3.282 | .031 |
| Cingulate_Post_R | Thal_MDl_R | 3.278 | .031 |
| Thal_VA_R | Thal_VL_L | 3.276 | .031 |
| Thal_LP_L | Thal_MDl_R | 3.271 | .031 |
| Cerebellum_Crus1_R | ACC_pre_L | 3.267 | .031 |
| OFClat_L | Thal_PuI_R | 3.256 | .031 |
| Thal_VPL_L | ACC_sub_L | 3.254 | .031 |
| Thal_LP_R | ACC_sub_R | 3.252 | .031 |
| Rectus_R | Pallidum_R | 3.251 | .031 |
| Thal_VPL_L | Thal_PuL_L | 3.249 | .031 |
| Lingual_L | Thal_LP_R | 3.248 | .031 |
| Cingulate_Post_L | Pallidum_R | 3.247 | .031 |
| Cerebellum_Crus2_L | Thal_VL_L | 3.243 | .031 |
| Cingulate_Post_L | Pallidum_L | 3.240 | .031 |
| Thal_PuI_R | N_Acc_L | 3.238 | .031 |
| Thal_LP_R | N_Acc_R | 3.234 | .031 |
| Cingulate_Post_R | Thal_LP_R | 3.234 | .031 |
| Thal_AV_L | Thal_PuA_R | 3.233 | .031 |
| Thal_PuM_R | Thal_PuL_L | 3.233 | .031 |
| Pallidum_L | Thal_MDl_R | 3.231 | .030 |
| Thal_MGN_L | Thal_PuA_L | 3.223 | .030 |
| Frontal_Sup_Medial_R | Cerebellum_4_5_R | 3.221 | .030 |
| Cingulate_Post_R | Pallidum_R | 3.220 | .030 |
| Thal_AV_L | Thal_MDm_L | 3.217 | .030 |
| Thal_VA_R | Thal_MDm_R | 3.216 | .030 |
| Thal_AV_L | Thal_MDl_L | 3.216 | .030 |
| Cingulate_Mid_R | Thal_LP_R | 3.209 | .030 |
| Thal_LP_R | Thal_PuM_R | 3.209 | .030 |
| Thal_AV_L | Thal_VL_L | 3.207 | .030 |
| OFCpost_L | Angular_L | 3.207 | .030 |
| Frontal_Med_Orb_L | Thal_LGN_L | 3.203 | .030 |
| Cerebellum_Crus2_R | Thal_LGN_L | 3.201 | .030 |
| Thal_LP_R | ACC_sup_R | 3.201 | .030 |
| Thal_MGN_L | Thal_PuI_R | 3.196 | .030 |
| Thal_MDl_L | Thal_LGN_L | 3.194 | .030 |
| Thal_LP_L | N_Acc_L | 3.185 | .030 |
| Thal_VL_R | Thal_MDl_L | 3.185 | .030 |
| Thal_VL_L | ACC_pre_L | 3.175 | .029 |
| Pallidum_L | Cerebellum_Crus2_L | 3.174 | .029 |
| Cingulate_Post_R | Vermis_9 | 3.173 | .029 |
| Thal_PuI_L | Thal_PuL_L | 3.164 | .029 |
| Cerebellum_Crus1_L | ACC_pre_L | 3.163 | .029 |
| Cingulate_Post_L | Thal_MDl_R | 3.160 | .029 |
| Thal_MDl_R | ACC_pre_L | 3.153 | .029 |
| Cingulate_Post_L | N_Acc_L | 3.153 | .029 |
| Temporal_Pole_Sup_L | Thal_PuI_R | 3.149 | .029 |
| Temporal_Pole_Mid_L | Thal_LP_R | 3.148 | .029 |
| Cerebellum_Crus2_R | Thal_VA_R | 3.148 | .029 |
| OFCant_L | Thal_VA_R | 3.145 | .029 |
| OFCmed_L | Thal_PuL_R | 3.145 | .029 |
| Heschl_R | Thal_PuA_L | 3.143 | .029 |
| Frontal_Sup_Medial_L | Pallidum_L | 3.140 | .029 |
| Thal_VL_L | Thal_LGN_L | 3.137 | .029 |
| Thal_Re_L | VTA_R | 3.137 | .029 |
| Thal_VPL_R | Thal_PuA_L | 3.134 | .029 |
| Pallidum_R | Thal_LP_L | 3.133 | .029 |
| Pallidum_L | Thal_LP_L | 3.132 | .029 |
| Putamen_L | Thal_PuM_L | 3.132 | .029 |
| VTA_L | SN_pc_L | 3.132 | .029 |
| Putamen_R | Pallidum_R | 3.131 | .029 |
| Cingulate_Post_L | Lingual_L | 3.129 | .029 |
| Hippocampus_R | Angular_L | 3.128 | .029 |
| Thal_LP_L | LC_L | 3.120 | .028 |
| Thal_PuA_R | LC_L | 3.120 | .028 |
| Thal_AV_L | Thal_VL_R | 3.107 | .028 |
| Putamen_L | Thal_PuA_L | 3.106 | .028 |
| Pallidum_L | Thal_PuM_L | 3.103 | .028 |
| Frontal_Sup_Medial_L | Thal_LP_R | 3.102 | .028 |
| Thal_PuA_R | Thal_PuL_L | 3.102 | .028 |
| Frontal_Sup_Medial_R | Pallidum_L | 3.101 | .028 |
| Thal_AV_L | ACC_sub_L | 3.101 | .028 |
| **Note:** AAL: Automated Anatomical Parcellation Atlas 3v2. The initial component-forming threshold was set to *t*<-3.1*.* | | | |

| **Supplementary Table 6.** Nodal degree and nodal strength of brain regions within a network, negatively associated with acute cortisol increase, identified by NBS. | | | |
| --- | --- | --- | --- |
| **AAL Index** | **Label** | **Nodal Degree** | **Nodal Strength** |
| 142 | Thal_PuA_L | 28 | 1.394 |
| 133 | Thal_MDl_R | 26 | 1.262 |
| 119 | Thal_LP_R | 29 | 1.191 |
| 130 | Thal_MDm_L | 18 | 1.013 |
| 140 | Thal_PuM_L | 21 | .934 |
| 141 | Thal_PuM_R | 20 | .917 |
| 122 | Thal_VL_L | 20 | .816 |
| 124 | Thal_VPL_L | 18 | .816 |
| 77 | Pallidum_R | 20 | .779 |
| 76 | Pallidum_L | 21 | .758 |
| 132 | Thal_MDl_L | 16 | .698 |
| 117 | Thal_AV_R | 15 | .689 |
| 143 | Thal_PuA_R | 15 | .662 |
| 37 | Cingulate_Post_R | 17 | .66 |
| 131 | Thal_MDm_R | 13 | .632 |
| 121 | Thal_VA_R | 15 | .573 |
| 123 | Thal_VL_R | 12 | .544 |
| 36 | Cingulate_Post_L | 12 | .415 |
| 116 | Thal_AV_L | 11 | .383 |
| 139 | Thal_PuI_R | 10 | .357 |
| 134 | Thal_LGN_L | 9 | .339 |
| 144 | Thal_PuL_L | 9 | .319 |
| 125 | Thal_VPL_R | 8 | .305 |
| 91 | Cerebellum_Crus1_R | 8 | .288 |
| 148 | ACC_pre_L | 8 | .271 |
| 152 | N_Acc_L | 7 | .247 |
| 145 | Thal_PuL_R | 5 | .216 |
| 120 | Thal_VA_L | 4 | .205 |
| 154 | VTA_L | 5 | .183 |
| 20 | Frontal_Med_Orb_L | 4 | .159 |
| 118 | Thal_LP_L | 5 | .147 |
| 74 | Putamen_L | 4 | .137 |
| 93 | Cerebellum_Crus2_R | 4 | .136 |
| 136 | Thal_MGN_L | 4 | .131 |
| 66 | Angular_L | 4 | .126 |
| 147 | ACC_sub_R | 3 | .112 |
| 114 | Vermis_9 | 3 | .107 |
| 162 | LC_L | 3 | .106 |
| 90 | Cerebellum_Crus1_L | 3 | .099 |
| 75 | Putamen_R | 3 | .097 |
| 48 | Lingual_L | 3 | .091 |
| 126 | Thal_IL_L | 2 | .089 |
| 127 | Thal_IL_R | 2 | .078 |
| 86 | Temporal_Pole_Mid_L | 2 | .073 |
| 79 | Heschl_R | 2 | .068 |
| 128 | Thal_Re_L | 2 | .067 |
| 138 | Thal_PuI_L | 2 | .065 |
| 99 | Cerebellum_6_R | 2 | .065 |
| 97 | Cerebellum_4_5_R | 2 | .064 |
| 23 | Rectus_R | 2 | .064 |
| 156 | SN_pc_L | 2 | .063 |
| 155 | VTA_R | 2 | .062 |
| 151 | ACC_sup_R | 2 | .062 |
| 92 | Cerebellum_Crus2_L | 2 | .06 |
| 146 | ACC_sub_L | 2 | .059 |
| 19 | Frontal_Sup_Medial_R | 2 | .058 |
| 40 | ParaHippocampal_L | 1 | .058 |
| 18 | Frontal_Sup_Medial_L | 2 | .057 |
| 129 | Thal_Re_R | 1 | .039 |
| 45 | Calcarine_R | 1 | .039 |
| 21 | Frontal_Med_Orb_R | 1 | .038 |
| 108 | Vermis_1_2 | 1 | .038 |
| 163 | LC_R | 1 | .036 |
| 22 | Rectus_L | 1 | .035 |
| 150 | ACC_sup_L | 1 | .035 |
| 110 | Vermis_4_5 | 1 | .035 |
| 44 | Calcarine_L | 1 | .034 |
| 135 | Thal_LGN_R | 1 | .033 |
| 43 | Amygdala_R | 1 | .032 |
| 30 | OFClat_L | 1 | .031 |
| 153 | N_Acc_R | 1 | .031 |
| 149 | ACC_pre_R | 1 | .031 |
| 28 | OFCpost_L | 1 | .03 |
| 35 | Cingulate_Mid_R | 1 | .03 |
| 24 | OFCmed_L | 1 | .029 |
| 26 | OFCant_L | 1 | .029 |
| 39 | Hippocampus_R | 1 | .029 |
| 82 | Temporal_Pole_Sup_L | 1 | .029 |
| **Note:** AAL: Automated Anatomical Parcellation Atlas 3v2. The initial component-forming threshold was set to *t*<-3.1*.* The effect size of the connections was used to compute nodal strength. | | | |

| **Supplementary Table 7.** Results of repeated NBS runs with various *t* thresholds. | | | | | |
| --- | --- | --- | --- | --- | --- |
| ***t* threshold** | ***p*** | ***p_FDR_*** | **No. Nodes** | **No. Edges** | ***µ*** $\boldsymbol{\eta}_{\boldsymbol{partial}}^{\boldsymbol{2}}$ |
| -2.0 | .022 | .045 | 164 | 1564 | .021 |
| -2.5 | .011 | .026 | 132 | 714 | .029 |
| -3.0 | .005 | .015 | 91 | 311 | .039 |
| -3.1 | .005 | .015 | 78 | 258 | .041 |
| -3.5 | .001 | .008 | 48 | 138 | .049 |
| -4.0 | .000 | .002 | 29 | 67 | .059 |
| **Note:** *t* < -3.1 has been selected as the threshold. The remaining work for the sensitivity analyses has been completed. Effect size was computed using partial eta squared ($\eta_{partial}^{2}$). FDR correction was performed across 12 t‑thresholds (6 positive, 6 negative). No significant results were found for any positive associations. | | | | | |

| **Supplementary Table 8.** Results of sex-specific NBS analyses. | | | | | |
| --- | --- | --- | --- | --- | --- |
| **Contrast** | ***p*** | ***p_FDR_*** | **No. Nodes** | **No. Edges** | ***µ*** $\boldsymbol{\eta}_{\boldsymbol{partial}}^{\boldsymbol{2}}$ |
| Male Negative | .033 | .074 | 53 | 65 | .035 |
| **Female Negative** | **.002** | **.011** | **112** | **337** | **.039** |
| Male Positive | .152 | .183 | - | - | - |
| Female Positive | .287 | .287 | - | - | - |
| Male > Female | .045 | .074 | 57 | 70 | .037 |
| Female > Male | .050 | .074 | 48 | 51 | .036 |
| **Note:** *t*<-3.1 has been selected as the threshold. Negative and positive models reveal negative and positive rsFC‑cortisol associations. Effect size was computed using partial eta squared ($\eta_{partial}^{2}$). FDR correction was performed across 6 contrasts. Significant results after FDR corrected are highlighted. | | | | | |

| **Supplementary Table 9.** Connections within a significant subnetwork in females, negatively associated with acute cortisol increase, as identified by NBS. | | | |
| --- | --- | --- | --- |
| **Region A** | **Region B** | ***t*** | $\boldsymbol{\eta}_{\boldsymbol{partial}}^{\boldsymbol{2}}$ |
| ParaHippocampal_L | Thal_LP_R | 6.678 | .119 |
| Thal_MDm_L | Thal_MDl_R | 5.462 | .083 |
| Thal_VA_L | Thal_PuA_L | 5.201 | .076 |
| Thal_LP_R | Thal_MDm_L | 5.189 | .075 |
| Cingulate_Post_R | Thal_PuM_R | 5.157 | .074 |
| Thal_AV_R | Thal_MDm_L | 5.113 | .073 |
| Thal_VPL_L | Thal_MDm_R | 4.938 | .069 |
| Thal_VPL_L | Thal_MDm_L | 4.878 | .067 |
| Thal_VL_L | Thal_MDm_L | 4.826 | .066 |
| Pallidum_L | Thal_MDm_L | 4.742 | .064 |
| Thal_VL_L | Thal_MDm_R | 4.711 | .063 |
| Thal_AV_R | Thal_VA_L | 4.697 | .062 |
| Thal_VL_L | Thal_MDl_R | 4.670 | .062 |
| Thal_LP_R | Thal_VA_L | 4.649 | .061 |
| Thal_MDl_R | Thal_PuM_R | 4.617 | .061 |
| Thal_VPL_L | Thal_MDl_R | 4.610 | .060 |
| Thal_MDm_L | Thal_MDm_R | 4.588 | .060 |
| Thal_MDm_R | Thal_MDl_L | 4.587 | .060 |
| Thal_AV_R | Thal_VPL_L | 4.573 | .059 |
| Thal_VL_L | Thal_PuM_R | 4.541 | .059 |
| Thal_VPL_L | Thal_PuM_R | 4.482 | .057 |
| Thal_AV_R | Thal_MDl_R | 4.475 | .057 |
| Thal_PuM_R | Thal_PuL_R | 4.461 | .057 |
| Rectus_R | Raphe_D | 4.424 | .056 |
| Thal_MDm_R | Thal_PuA_L | 4.419 | .056 |
| Frontal_Med_Orb_R | Raphe_D | 4.414 | .056 |
| Thal_VA_L | Thal_MDl_L | 4.412 | .056 |
| Thal_AV_L | Thal_PuM_R | 4.395 | .055 |
| Pallidum_L | Thal_VPL_R | 4.386 | .055 |
| Cingulate_Post_R | Thal_VL_L | 4.376 | .055 |
| Thal_AV_R | Thal_PuM_R | 4.371 | .055 |
| Thal_MDl_R | Thal_PuA_R | 4.370 | .055 |
| Thal_MDm_L | Thal_PuM_L | 4.361 | .054 |
| Thal_MDl_R | Thal_PuM_L | 4.309 | .053 |
| Thal_VL_L | Thal_MDl_L | 4.301 | .053 |
| Thal_LP_R | Thal_MDl_R | 4.300 | .053 |
| Olfactory_R | Raphe_D | 4.281 | .052 |
| Cerebellum_4_5_R | Thal_LP_R | 4.280 | .052 |
| Thal_AV_R | Thal_PuM_L | 4.278 | .052 |
| Cingulate_Post_R | Thal_AV_R | 4.263 | .052 |
| Thal_MDm_L | Thal_PuM_R | 4.249 | .052 |
| Thal_MGN_R | SN_pr_R | 4.200 | .051 |
| Pallidum_R | Thal_PuI_R | 4.191 | .050 |
| Thal_AV_R | Thal_VL_R | 4.188 | .050 |
| Pallidum_R | Thal_LGN_R | 4.187 | .050 |
| Thal_VPL_L | Thal_PuM_L | 4.155 | .050 |
| Pallidum_L | Thal_VL_R | 4.149 | .049 |
| Thal_LP_R | Thal_PuM_R | 4.122 | .049 |
| Thal_LP_L | SN_pr_R | 4.117 | .049 |
| Cerebellum_Crus1_R | Thal_LP_R | 4.108 | .049 |
| Cingulate_Post_R | Thal_LP_R | 4.090 | .048 |
| Thal_VPL_R | SN_pr_R | 4.087 | .048 |
| Thal_VA_R | Thal_MDm_L | 4.085 | .048 |
| Thal_LP_R | Thal_VPL_R | 4.081 | .048 |
| Thal_VPL_L | Thal_PuA_R | 4.081 | .048 |
| Pallidum_L | Thal_VL_L | 4.070 | .048 |
| Thal_AV_R | Thal_PuA_L | 4.064 | .048 |
| Thal_MDl_L | Thal_MDl_R | 4.058 | .047 |
| Pallidum_L | Thal_MDl_R | 4.053 | .047 |
| Pallidum_L | Thal_MDl_L | 4.049 | .047 |
| Thal_VL_R | Thal_PuM_L | 4.048 | .047 |
| Thal_LP_R | Thal_PuA_R | 4.047 | .047 |
| Thal_LP_R | N_Acc_R | 4.045 | .047 |
| Thal_VPL_L | SN_pr_R | 4.037 | .047 |
| Vermis_4_5 | Thal_PuI_R | 4.037 | .047 |
| Calcarine_R | Thal_PuI_R | 4.034 | .047 |
| Thal_PuA_R | Thal_PuL_R | 4.030 | .047 |
| Cerebellum_6_R | Thal_LP_R | 4.020 | .047 |
| Cingulate_Mid_L | Thal_LP_R | 4.018 | .047 |
| Frontal_Med_Orb_L | Thal_LP_R | 4.004 | .046 |
| Thal_VA_L | Thal_VA_R | 4.000 | .046 |
| Thal_VL_L | Thal_VPL_L | 3.998 | .046 |
| Pallidum_L | Cerebellum_Crus2_R | 3.986 | .046 |
| Thal_MDm_L | Thal_PuA_R | 3.985 | .046 |
| Thal_VA_L | Thal_PuM_R | 3.975 | .046 |
| Vermis_4_5 | SN_pc_R | 3.966 | .045 |
| ParaHippocampal_L | Thal_LP_L | 3.964 | .045 |
| Cingulate_Post_L | Thal_PuA_L | 3.945 | .045 |
| Thal_MDl_L | Thal_PuM_L | 3.922 | .044 |
| Precuneus_L | Thal_LP_R | 3.916 | .044 |
| Thal_VA_L | Thal_PuM_L | 3.905 | .044 |
| Angular_R | Raphe_D | 3.903 | .044 |
| Thal_VL_R | Thal_MDm_L | 3.893 | .044 |
| Angular_R | Thal_PuM_R | 3.892 | .044 |
| Thal_VL_L | Thal_PuA_L | 3.885 | .044 |
| Thal_PuI_L | VTA_R | 3.884 | .044 |
| Olfactory_L | Raphe_D | 3.879 | .043 |
| Thal_AV_R | Thal_VL_L | 3.875 | .043 |
| Thal_LP_R | Thal_MDm_R | 3.874 | .043 |
| Thal_VA_R | Thal_PuM_R | 3.872 | .043 |
| Hippocampus_R | Angular_L | 3.862 | .043 |
| Pallidum_L | Cerebellum_Crus1_R | 3.859 | .043 |
| Vermis_4_5 | Thal_VL_L | 3.857 | .043 |
| Thal_VL_L | Thal_VL_R | 3.857 | .043 |
| Thal_VA_L | Thal_MDm_R | 3.854 | .043 |
| Temporal_Inf_R | Thal_LP_R | 3.854 | .043 |
| Thal_MDl_L | SN_pr_R | 3.854 | .043 |
| OFCant_L | Thal_LP_R | 3.846 | .043 |
| ACC_pre_R | Raphe_D | 3.845 | .043 |
| Thal_PuA_L | ACC_pre_L | 3.844 | .043 |
| Cingulate_Post_L | SN_pc_L | 3.836 | .043 |
| Rectus_L | Raphe_D | 3.836 | .043 |
| Thal_LP_R | Thal_MDl_L | 3.827 | .042 |
| Cerebellum_6_R | Thal_VL_L | 3.826 | .042 |
| Thal_PuA_L | Thal_PuA_R | 3.826 | .042 |
| Thal_AV_R | Thal_PuA_R | 3.826 | .042 |
| Cingulate_Mid_R | Thal_LP_R | 3.822 | .042 |
| Cerebellum_Crus1_R | Thal_PuA_L | 3.812 | .042 |
| Cingulate_Post_R | Thal_MDm_L | 3.800 | .042 |
| Thal_LP_R | ACC_pre_L | 3.796 | .042 |
| Thal_MDl_L | Thal_PuA_R | 3.795 | .042 |
| Cingulate_Post_L | Thal_AV_R | 3.795 | .042 |
| Thal_VL_R | Thal_PuA_R | 3.794 | .042 |
| Thal_VA_R | Thal_PuL_R | 3.789 | .042 |
| Thal_MDl_L | Thal_PuM_R | 3.788 | .042 |
| Thal_AV_R | Thal_MDl_L | 3.762 | .041 |
| Temporal_Pole_Mid_L | Thal_LP_R | 3.757 | .041 |
| Thal_VPL_R | Thal_PuM_R | 3.755 | .041 |
| Pallidum_R | Thal_MDl_L | 3.753 | .041 |
| Thal_VL_L | SN_pr_L | 3.751 | .041 |
| Rectus_L | SN_pr_L | 3.751 | .041 |
| Frontal_Med_Orb_L | VTA_L | 3.748 | .041 |
| ACC_sup_R | Raphe_D | 3.738 | .040 |
| Cingulate_Post_L | ParaHippocampal_L | 3.727 | .040 |
| Thal_MDm_R | Thal_PuM_R | 3.727 | .040 |
| Thal_AV_R | Thal_MDm_R | 3.725 | .040 |
| Thal_VA_L | Thal_VL_L | 3.705 | .040 |
| Thal_VA_L | Thal_VL_R | 3.695 | .040 |
| VTA_L | SN_pc_L | 3.695 | .040 |
| Thal_LP_R | ACC_sup_R | 3.690 | .040 |
| Cerebellum_Crus1_R | Thal_MDm_L | 3.682 | .039 |
| Frontal_Sup_2_R | Thal_PuM_R | 3.681 | .039 |
| Thal_PuI_R | Thal_PuM_R | 3.681 | .039 |
| SN_pr_L | SN_pr_R | 3.672 | .039 |
| Cerebellum_Crus2_R | Thal_PuA_L | 3.671 | .039 |
| Thal_LP_R | Thal_PuM_L | 3.669 | .039 |
| Thal_MDm_L | Thal_PuA_L | 3.669 | .039 |
| Thal_PuL_L | SN_pr_R | 3.666 | .039 |
| Thal_LP_L | N_Acc_L | 3.665 | .039 |
| Cingulate_Post_R | SN_pc_L | 3.664 | .039 |
| Rectus_L | VTA_R | 3.664 | .039 |
| Thal_AV_R | ACC_pre_L | 3.647 | .039 |
| Cingulate_Post_L | Thal_PuM_R | 3.644 | .039 |
| Thal_VL_R | Thal_MDl_R | 3.641 | .039 |
| Pallidum_L | N_Acc_L | 3.628 | .038 |
| Vermis_6 | Thal_PuI_R | 3.626 | .038 |
| Thal_LP_R | N_Acc_L | 3.623 | .038 |
| Temporal_Pole_Mid_L | Thal_PuL_L | 3.618 | .038 |
| SupraMarginal_L | Thal_LP_R | 3.609 | .038 |
| Pallidum_L | Cerebellum_6_R | 3.607 | .038 |
| Cingulate_Post_L | Thal_VPL_L | 3.606 | .038 |
| Thal_MDl_R | SN_pc_R | 3.605 | .038 |
| Pallidum_L | Thal_IL_L | 3.601 | .038 |
| Cingulate_Post_L | Cerebellum_6_R | 3.592 | .038 |
| Thal_LP_R | ACC_pre_R | 3.589 | .037 |
| Thal_MGN_R | Raphe_M | 3.589 | .037 |
| Thal_VPL_R | Thal_MDm_R | 3.584 | .037 |
| Cerebellum_4_5_R | Thal_LGN_R | 3.576 | .037 |
| Thal_Re_L | VTA_R | 3.575 | .037 |
| Parietal_Inf_L | Raphe_M | 3.574 | .037 |
| Thal_MDl_R | ACC_pre_L | 3.574 | .037 |
| Cerebellum_Crus1_L | Thal_LP_R | 3.574 | .037 |
| Cingulate_Post_R | Vermis_4_5 | 3.572 | .037 |
| Calcarine_L | Thal_PuI_R | 3.572 | .037 |
| SN_pc_L | SN_pr_R | 3.560 | .037 |
| Cerebellum_4_5_R | Thal_LP_L | 3.558 | .037 |
| Thal_MDl_L | Thal_PuA_L | 3.551 | .037 |
| Frontal_Sup_2_L | Thal_MDm_L | 3.549 | .037 |
| Hippocampus_L | Raphe_D | 3.547 | .037 |
| Thal_MDl_R | Thal_MGN_L | 3.542 | .037 |
| Frontal_Inf_Orb_2_R | Thal_MDl_R | 3.529 | .036 |
| Thal_PuA_R | ACC_sup_L | 3.528 | .036 |
| Hippocampus_L | Hippocampus_R | 3.521 | .036 |
| Cerebellum_6_R | Thal_PuM_R | 3.521 | .036 |
| Putamen_R | Pallidum_L | 3.516 | .036 |
| Cerebellum_Crus1_R | Thal_PuI_R | 3.512 | .036 |
| Frontal_Sup_2_L | Thal_LP_R | 3.510 | .036 |
| Hippocampus_R | Thal_PuL_L | 3.506 | .036 |
| Thal_VL_L | Thal_IL_R | 3.501 | .036 |
| Cerebellum_4_5_R | SN_pc_R | 3.497 | .036 |
| Thal_VL_L | Thal_PuA_R | 3.496 | .036 |
| Thal_LP_R | Thal_VPL_L | 3.490 | .035 |
| Frontal_Med_Orb_L | Raphe_D | 3.489 | .035 |
| Lingual_L | Thal_LP_R | 3.479 | .035 |
| Thal_PuM_R | SN_pc_L | 3.466 | .035 |
| Rectus_R | VTA_R | 3.458 | .035 |
| Cerebellum_Crus1_R | Thal_PuM_R | 3.458 | .035 |
| Thal_VA_R | Thal_PuM_L | 3.455 | .035 |
| Cerebellum_Crus2_R | Thal_VA_L | 3.453 | .035 |
| Thal_VPL_L | Thal_MDl_L | 3.452 | .035 |
| Lingual_L | Thal_PuI_R | 3.451 | .035 |
| Cerebellum_4_5_R | Thal_PuI_R | 3.450 | .035 |
| Cerebellum_Crus2_R | Thal_VA_R | 3.448 | .035 |
| Cerebellum_Crus1_R | Thal_VL_L | 3.444 | .035 |
| Thal_VL_R | Thal_PuA_L | 3.441 | .035 |
| Frontal_Sup_Medial_L | Thal_AV_R | 3.441 | .035 |
| Pallidum_L | Pallidum_R | 3.434 | .034 |
| Cingulate_Post_L | Cerebellum_Crus1_R | 3.431 | .034 |
| Frontal_Med_Orb_L | Pallidum_L | 3.429 | .034 |
| Thal_AV_L | Thal_PuA_L | 3.422 | .034 |
| Cerebellum_Crus1_L | Thal_PuM_R | 3.421 | .034 |
| Frontal_Med_Orb_R | Thal_LP_R | 3.418 | .034 |
| Pallidum_R | Thal_MDm_L | 3.417 | .034 |
| Vermis_4_5 | Thal_MDm_L | 3.417 | .034 |
| Cingulate_Post_R | Thal_MDl_L | 3.415 | .034 |
| Fusiform_L | Thal_LP_R | 3.412 | .034 |
| ParaHippocampal_L | Raphe_D | 3.412 | .034 |
| Thal_VL_L | Thal_PuM_L | 3.411 | .034 |
| Thal_LP_R | ACC_sup_L | 3.411 | .034 |
| Cingulate_Post_R | Thal_VPL_L | 3.410 | .034 |
| Thal_VL_R | Thal_MDl_L | 3.405 | .034 |
| Caudate_R | Thal_PuM_R | 3.403 | .034 |
| Heschl_R | Thal_AV_R | 3.403 | .034 |
| Thal_MDm_R | Thal_MDl_R | 3.402 | .034 |
| Thal_AV_L | Thal_MDl_L | 3.395 | .034 |
| Thal_VA_L | Thal_MDl_R | 3.385 | .033 |
| Frontal_Sup_Medial_L | Thal_LP_R | 3.384 | .033 |
| Vermis_6 | Thal_MDm_L | 3.383 | .033 |
| Cingulate_Mid_R | Thal_AV_R | 3.378 | .033 |
| Thal_VPL_L | Thal_PuL_R | 3.374 | .033 |
| Thal_AV_L | Thal_MDm_L | 3.373 | .033 |
| Thal_AV_L | Thal_VL_L | 3.372 | .033 |
| ACC_sub_R | VTA_L | 3.372 | .033 |
| Thal_MDl_R | Thal_PuI_R | 3.364 | .033 |
| Thal_MDl_R | ACC_sup_L | 3.363 | .033 |
| Thal_VA_L | Thal_MDm_L | 3.358 | .033 |
| Cingulate_Post_L | Cerebellum_4_5_R | 3.352 | .033 |
| Thal_LP_L | Thal_MDl_R | 3.349 | .033 |
| Pallidum_L | Thal_AV_R | 3.344 | .033 |
| Cerebellum_4_5_L | Thal_PuI_R | 3.342 | .033 |
| Pallidum_R | Thal_VL_L | 3.337 | .033 |
| ParaHippocampal_R | Thal_LP_R | 3.336 | .033 |
| Putamen_R | Pallidum_R | 3.330 | .032 |
| Thal_AV_R | Thal_PuL_L | 3.329 | .032 |
| Thal_MDm_L | Thal_PuL_L | 3.329 | .032 |
| Thal_LP_L | Thal_MDm_L | 3.328 | .032 |
| OFCpost_L | Thal_LP_L | 3.326 | .032 |
| Cingulate_Post_L | Thal_VL_L | 3.324 | .032 |
| Cerebellum_Crus2_L | Thal_VA_L | 3.321 | .032 |
| Cingulate_Post_R | Thal_LP_L | 3.321 | .032 |
| Thal_PuM_R | Thal_PuA_L | 3.319 | .032 |
| Thal_AV_L | Thal_PuM_L | 3.319 | .032 |
| Vermis_6 | Thal_LP_R | 3.317 | .032 |
| Cerebellum_Crus2_L | Thal_VL_L | 3.305 | .032 |
| Putamen_L | Thal_LP_R | 3.299 | .032 |
| Cingulate_Post_L | Hippocampus_R | 3.298 | .032 |
| Cerebellum_6_R | Thal_PuA_L | 3.297 | .032 |
| Cingulate_Post_L | Thal_MDm_L | 3.295 | .032 |
| Hippocampus_L | Thal_LP_R | 3.291 | .032 |
| Thal_VPL_L | Thal_MGN_L | 3.291 | .032 |
| SupraMarginal_L | Thal_AV_R | 3.287 | .032 |
| OFCpost_R | Raphe_D | 3.283 | .032 |
| Thal_Re_R | SN_pr_R | 3.280 | .031 |
| Thal_MDl_R | ACC_sup_R | 3.279 | .031 |
| Thal_MDm_L | ACC_pre_R | 3.273 | .031 |
| SN_pc_L | Raphe_D | 3.268 | .031 |
| Thal_VL_L | ACC_pre_L | 3.267 | .031 |
| Lingual_R | Thal_LP_R | 3.260 | .031 |
| SN_pc_L | SN_pc_R | 3.260 | .031 |
| Cerebellum_Crus1_L | Thal_PuA_L | 3.257 | .031 |
| Cingulate_Post_R | VTA_L | 3.255 | .031 |
| Temporal_Pole_Mid_R | Thal_LP_R | 3.251 | .031 |
| Frontal_Mid_2_L | Thal_MDl_R | 3.248 | .031 |
| Angular_R | Thal_LP_R | 3.246 | .031 |
| Pallidum_R | Thal_PuA_L | 3.246 | .031 |
| Pallidum_R | Thal_PuM_R | 3.245 | .031 |
| Pallidum_R | Thal_MDl_R | 3.238 | .031 |
| Cerebellum_Crus1_R | Thal_MDl_L | 3.237 | .031 |
| Thal_IL_R | Thal_MDl_R | 3.233 | .031 |
| Temporal_Pole_Mid_L | Thal_LP_L | 3.233 | .031 |
| SN_pr_R | Red_N_R | 3.229 | .031 |
| Pallidum_R | Thal_IL_L | 3.228 | .031 |
| Thal_VPL_R | Thal_MDl_R | 3.227 | .030 |
| Thal_MDm_L | ACC_sup_R | 3.223 | .030 |
| OFClat_R | LC_R | 3.219 | .030 |
| Pallidum_L | Thal_VPL_L | 3.214 | .030 |
| Heschl_R | Thal_PuA_L | 3.212 | .030 |
| LC_R | Raphe_D | 3.212 | .030 |
| Thal_IL_R | SN_pc_R | 3.211 | .030 |
| Rectus_L | VTA_L | 3.209 | .030 |
| Thal_VL_R | Thal_PuM_R | 3.205 | .030 |
| Parietal_Inf_R | Thal_MDl_R | 3.200 | .030 |
| Cerebellum_8_R | Thal_LP_R | 3.200 | .030 |
| OFCmed_R | Red_N_R | 3.199 | .030 |
| Cingulate_Post_L | VTA_L | 3.199 | .030 |
| Pallidum_R | Thal_LP_L | 3.198 | .030 |
| Precuneus_L | Raphe_D | 3.197 | .030 |
| Cuneus_L | Thal_PuI_R | 3.197 | .030 |
| Thal_PuL_R | ACC_pre_L | 3.195 | .030 |
| Heschl_R | Thal_LP_R | 3.193 | .030 |
| Thal_VA_R | Thal_MDl_L | 3.192 | .030 |
| Supp_Motor_Area_L | Thal_LP_R | 3.192 | .030 |
| Insula_R | SN_pc_R | 3.190 | .030 |
| OFCpost_L | Thal_LP_R | 3.190 | .030 |
| Thal_LP_R | Thal_PuA_L | 3.190 | .030 |
| Frontal_Sup_Medial_R | Thal_LP_R | 3.188 | .030 |
| Thal_AV_L | Thal_MDm_R | 3.186 | .030 |
| Cerebellum_4_5_L | Thal_LP_R | 3.185 | .030 |
| Putamen_R | Thal_LP_R | 3.183 | .030 |
| Thal_PuM_L | Thal_PuL_R | 3.183 | .030 |
| Cerebellum_6_R | Thal_PuI_R | 3.177 | .030 |
| Thal_LP_R | Thal_VL_R | 3.177 | .030 |
| Thal_VL_L | SN_pr_R | 3.172 | .030 |
| Pallidum_L | Temporal_Pole_Mid_L | 3.167 | .029 |
| Caudate_R | Thal_PuI_R | 3.161 | .029 |
| Frontal_Sup_Medial_R | Thal_PuM_R | 3.160 | .029 |
| Angular_L | Cerebellum_4_5_R | 3.151 | .029 |
| Thal_PuM_L | Thal_PuM_R | 3.149 | .029 |
| Thal_LP_R | LC_R | 3.149 | .029 |
| Cerebellum_9_L | Thal_LP_R | 3.147 | .029 |
| Thal_AV_R | Thal_VA_R | 3.146 | .029 |
| Cerebellum_Crus1_R | Thal_VA_L | 3.145 | .029 |
| Thal_MDl_R | VTA_L | 3.143 | .029 |
| Pallidum_L | Vermis_6 | 3.140 | .029 |
| Cerebellum_6_L | Thal_LP_R | 3.139 | .029 |
| Thal_MDl_R | Thal_PuA_L | 3.137 | .029 |
| Temporal_Pole_Mid_R | Raphe_D | 3.135 | .029 |
| Frontal_Sup_2_R | Raphe_D | 3.134 | .029 |
| OFClat_L | Thal_PuI_R | 3.133 | .029 |
| Pallidum_L | Temporal_Mid_R | 3.131 | .029 |
| VTA_L | Red_N_L | 3.131 | .029 |
| Pallidum_R | Thal_LGN_L | 3.130 | .029 |
| Fusiform_R | Thal_LP_R | 3.125 | .029 |
| Frontal_Sup_Medial_L | Pallidum_L | 3.121 | .029 |
| Cerebellum_4_5_L | SN_pc_R | 3.114 | .028 |
| Cerebellum_Crus1_L | Thal_VA_L | 3.109 | .028 |
| OFCpost_R | Thal_LP_R | 3.108 | .028 |
| Thal_MDl_L | ACC_pre_L | 3.107 | .028 |
| Frontal_Mid_2_L | Thal_MDm_L | 3.107 | .028 |
| Temporal_Pole_Mid_L | Thal_AV_L | 3.106 | .028 |
| Cingulate_Post_L | Vermis_4_5 | 3.106 | .028 |
| N_Acc_R | Raphe_D | 3.101 | .028 |
| Thal_LP_R | Thal_IL_R | 3.101 | .028 |
| Frontal_Mid_2_L | Thal_PuM_R | 3.101 | .028 |
| Thal_VPL_L | Thal_PuA_L | 3.101 | .028 |
| Thal_VPL_L | Thal_VPL_R | 3.100 | .028 |
| Precuneus_R | Raphe_D | 3.100 | .028 |
| **Note:** AAL: Automated Anatomical Parcellation Atlas 3v2. The initial component-forming threshold was set to *t*<-3.1*.* | | | |

| **Supplementary Table 10.** The nodal degree and nodal strength of brain regions within a significant subnetwork in females, negatively associated with acute cortisol increase, as identified by NBS. | | | |
| --- | --- | --- | --- |
| **AAL Index** | **Label** | **Nodal Degree** | **Nodal Strength** |
| 119 | Thal_LP_R | 57 | .345 |
| 141 | Thal_PuM_R | 29 | .176 |
| 133 | Thal_MDl_R | 28 | .170 |
| 130 | Thal_MDm_L | 27 | .164 |
| 122 | Thal_VL_L | 25 | .152 |
| 117 | Thal_AV_R | 22 | .133 |
| 142 | Thal_PuA_L | 21 | .127 |
| 132 | Thal_MDl_L | 20 | .121 |
| 76 | Pallidum_L | 20 | .121 |
| 164 | Raphe_D | 19 | .115 |
| 124 | Thal_VPL_L | 18 | .109 |
| 120 | Thal_VA_L | 16 | .097 |
| 139 | Thal_PuI_R | 15 | .091 |
| 36 | Cingulate_Post_L | 14 | .085 |
| 140 | Thal_PuM_L | 13 | .079 |
| 77 | Pallidum_R | 13 | .079 |
| 123 | Thal_VL_R | 12 | .073 |
| 131 | Thal_MDm_R | 12 | .073 |
| 37 | Cingulate_Post_R | 11 | .067 |
| 159 | SN_pr_R | 11 | .067 |
| 143 | Thal_PuA_R | 11 | .067 |
| 118 | Thal_LP_L | 10 | .061 |
| 91 | Cerebellum_Crus1_R | 10 | .061 |
| 121 | Thal_VA_R | 8 | .048 |
| 154 | VTA_L | 8 | .048 |
| 116 | Thal_AV_L | 8 | .048 |
| 97 | Cerebellum_4_5_R | 7 | .042 |
| 148 | ACC_pre_L | 7 | .042 |
| 156 | SN_pc_L | 7 | .042 |
| 125 | Thal_VPL_R | 7 | .042 |
| 157 | SN_pc_R | 7 | .042 |
| 99 | Cerebellum_6_R | 7 | .042 |
| 110 | Vermis_4_5 | 6 | .036 |
| 145 | Thal_PuL_R | 6 | .036 |
| 144 | Thal_PuL_L | 5 | .030 |
| 86 | Temporal_Pole_Mid_L | 5 | .030 |
| 127 | Thal_IL_R | 4 | .024 |
| 93 | Cerebellum_Crus2_R | 4 | .024 |
| 90 | Cerebellum_Crus1_L | 4 | .024 |
| 111 | Vermis_6 | 4 | .024 |
| 151 | ACC_sup_R | 4 | .024 |
| 40 | ParaHippocampal_L | 4 | .024 |
| 20 | Frontal_Med_Orb_L | 4 | .024 |
| 39 | Hippocampus_R | 4 | .024 |
| 22 | Rectus_L | 4 | .024 |
| 155 | VTA_R | 4 | .024 |
| 152 | N_Acc_L | 3 | .018 |
| 4 | Frontal_Mid_2_L | 3 | .018 |
| 163 | LC_R | 3 | .018 |
| 18 | Frontal_Sup_Medial_L | 3 | .018 |
| 158 | SN_pr_L | 3 | .018 |
| 67 | Angular_R | 3 | .018 |
| 150 | ACC_sup_L | 3 | .018 |
| 149 | ACC_pre_R | 3 | .018 |
| 38 | Hippocampus_L | 3 | .018 |
| 79 | Heschl_R | 3 | .018 |
| 75 | Putamen_R | 3 | .018 |
| 96 | Cerebellum_4_5_L | 3 | .018 |
| 135 | Thal_LGN_R | 2 | .012 |
| 153 | N_Acc_R | 2 | .012 |
| 137 | Thal_MGN_R | 2 | .012 |
| 161 | Red_N_R | 2 | .012 |
| 136 | Thal_MGN_L | 2 | .012 |
| 126 | Thal_IL_L | 2 | .012 |
| 2 | Frontal_Sup_2_L | 2 | .012 |
| 48 | Lingual_L | 2 | .012 |
| 68 | Precuneus_L | 2 | .012 |
| 19 | Frontal_Sup_Medial_R | 2 | .012 |
| 21 | Frontal_Med_Orb_R | 2 | .012 |
| 23 | Rectus_R | 2 | .012 |
| 28 | OFCpost_L | 2 | .012 |
| 29 | OFCpost_R | 2 | .012 |
| 35 | Cingulate_Mid_R | 2 | .012 |
| 64 | SupraMarginal_L | 2 | .012 |
| 66 | Angular_L | 2 | .012 |
| 165 | Raphe_M | 2 | .012 |
| 3 | Frontal_Sup_2_R | 2 | .012 |
| 92 | Cerebellum_Crus2_L | 2 | .012 |
| 73 | Caudate_R | 2 | .012 |
| 87 | Temporal_Pole_Mid_R | 2 | .012 |
| 31 | OFClat_R | 1 | .006 |
| 30 | OFClat_L | 1 | .006 |
| 89 | Temporal_Inf_R | 1 | .006 |
| 26 | OFCant_L | 1 | .006 |
| 25 | OFCmed_R | 1 | .006 |
| 128 | Thal_Re_L | 1 | .006 |
| 34 | Cingulate_Mid_L | 1 | .006 |
| 98 | Cerebellum_6_L | 1 | .006 |
| 17 | Olfactory_R | 1 | .006 |
| 160 | Red_N_L | 1 | .006 |
| 16 | Olfactory_L | 1 | .006 |
| 14 | Supp_Motor_Area_L | 1 | .006 |
| 11 | Frontal_Inf_Orb_2_R | 1 | .006 |
| 33 | Insula_R | 1 | .006 |
| 147 | ACC_sub_R | 1 | .006 |
| 129 | Thal_Re_R | 1 | .006 |
| 56 | Fusiform_L | 1 | .006 |
| 69 | Precuneus_R | 1 | .006 |
| 74 | Putamen_L | 1 | .006 |
| 63 | Parietal_Inf_R | 1 | .006 |
| 134 | Thal_LGN_L | 1 | .006 |
| 62 | Parietal_Inf_L | 1 | .006 |
| 57 | Fusiform_R | 1 | .006 |
| 138 | Thal_PuI_L | 1 | .006 |
| 85 | Temporal_Mid_R | 1 | .006 |
| 49 | Lingual_R | 1 | .006 |
| 103 | Cerebellum_8_R | 1 | .006 |
| 46 | Cuneus_L | 1 | .006 |
| 45 | Calcarine_R | 1 | .006 |
| 44 | Calcarine_L | 1 | .006 |
| 41 | ParaHippocampal_R | 1 | .006 |
| 104 | Cerebellum_9_L | 1 | .006 |
| **Note:** AAL: Automated Anatomical Parcellation Atlas 3v2. The initial component-forming threshold was set to *t*<-3.1*.* The effect size of the connections was used to compute nodal strength. | | | |

| **Supplementary Table 11.** Nodal degree and nodal strength of brain regions within the most relevant network identified by NBS-Predict (weight threshold=1.0). | | |
| --- | --- | --- |
| **Label** | **Nodal Degree** | **Nodal Strength** |
| Thal_PuA_L | 9 | 9.000 |
| Thal_MDl_R | 8 | 8.028 |
| Thal_PuM_R | 5 | 5.036 |
| Thal_MDm_L | 5 | 5.000 |
| Thal_VPL_L | 4 | 4.000 |
| Thal_LP_R | 3 | 3.000 |
| Thal_VL_R | 3 | 3.020 |
| Thal_MDm_R | 3 | 3.000 |
| Thal_PuA_R | 2 | 2.000 |
| Thal_PuM_L | 2 | 2.000 |
| Thal_AV_R | 2 | 2.028 |
| Pallidum_L | 2 | 2.020 |
| Thal_MDl_L | 2 | 2.000 |
| Frontal_Med_Orb_L | 1 | 1.000 |
| Thal_VA_L | 1 | 1.000 |
| Thal_AV_L | 1 | 1.036 |
| Thal_VPL_R | 1 | 1.000 |

**Note:** Nodal strength was computed from the edge weight of connections linking the corresponding brain regions.

| **Supplementary Table 12.** Nodal degree and nodal strength of brain regions within a highly robust network identified by NBS-Predict (weight threshold≥.9). | | |
| --- | --- | --- |
| **Label** | **Nodal Degree** | **Nodal Strength** |
| Thal_MDl_R | 12 | 11.871 |
| Thal_PuA_L | 12 | 11.893 |
| Thal_MDm_L | 11 | 10.671 |
| Thal_PuM_R | 10 | 9.822 |
| Thal_PuM_L | 8 | 7.614 |
| Pallidum_R | 7 | 6.812 |
| Thal_MDl_L | 7 | 6.824 |
| Thal_LP_R | 7 | 6.781 |
| Thal_VPL_L | 7 | 6.751 |
| Thal_AV_R | 6 | 5.726 |
| Thal_VL_R | 6 | 5.857 |
| Thal_VL_L | 5 | 4.960 |
| Thal_MDm_R | 5 | 4.984 |
| Pallidum_L | 5 | 4.842 |
| Thal_PuA_R | 4 | 3.888 |
| Cingulate_Post_R | 4 | 3.807 |
| Thal_LGN_L | 2 | 1.943 |
| Thal_PuL_R | 2 | 1.891 |
| N_Acc_L | 1 | .914 |
| Frontal_Med_Orb_L | 1 | 1.000 |
| Thal_IL_L | 1 | .938 |
| Thal_PuI_R | 1 | 1.000 |
| Thal_VPL_R | 1 | 1.000 |
| Thal_VA_R | 1 | .947 |
| Thal_VA_L | 1 | 1.000 |
| Thal_AV_L | 1 | 1.036 |
| ParaHippocampal_L | 1 | .945 |
| LC_L | 1 | .949 |

**Note:** Nodal strength was computed from the edge weight of connections linking the corresponding brain regions.

| **Supplementary Table 13.** Connections present within robust connected network identified by NBS-Predict (weight threshold≥.9). | | |
| --- | --- | --- |
| **Region A** | **Region B** | **Weight** |
| Thal_AV_L | Thal_PuM_R | 1.000 |
| Pallidum_R | Thal_LGN_L | 1.000 |
| Thal_AV_R | Thal_MDl_R | 1.000 |
| Pallidum_L | Thal_VL_R | 1.000 |
| Frontal_Med_Orb_L | Pallidum_L | 1.000 |
| Pallidum_R | Thal_PuI_R | 1.000 |
| Thal_AV_R | Thal_PuA_L | 1.000 |
| Thal_LP_R | Thal_VPL_R | 1.000 |
| Thal_LP_R | Thal_MDm_L | 1.000 |
| Thal_LP_R | Thal_MDl_R | 1.000 |
| Thal_VA_L | Thal_PuA_L | 1.000 |
| Thal_VL_R | Thal_PuM_L | 1.000 |
| Thal_VL_R | Thal_PuA_L | 1.000 |
| Thal_VPL_L | Thal_MDm_L | 1.000 |
| Thal_VPL_L | Thal_MDm_R | 1.000 |
| Thal_VPL_L | Thal_MDl_R | 1.000 |
| Thal_VPL_L | Thal_PuM_R | 1.000 |
| Thal_MDm_L | Thal_MDm_R | 1.000 |
| Thal_MDm_L | Thal_MDl_R | 1.000 |
| Thal_MDm_L | Thal_PuA_L | 1.000 |
| Thal_MDm_R | Thal_PuA_L | 1.000 |
| Thal_MDl_L | Thal_PuM_R | 1.000 |
| Thal_MDl_L | Thal_PuA_L | 1.000 |
| Thal_MDl_R | Thal_PuM_L | 1.000 |
| Thal_MDl_R | Thal_PuM_R | 1.000 |
| Thal_MDl_R | Thal_PuA_L | 1.000 |
| Thal_MDl_R | Thal_PuA_R | 1.000 |
| Thal_PuM_R | Thal_PuA_L | 1.000 |
| Thal_PuA_L | Thal_PuA_R | 1.000 |
| Cingulate_Post_R | Thal_VL_L | .992 |
| Thal_VL_L | Thal_MDm_L | .992 |
| Thal_VL_L | Thal_MDm_R | .992 |
| Thal_VL_L | Thal_MDl_R | .992 |
| Thal_VL_L | Thal_PuA_L | .992 |
| Thal_MDm_R | Thal_MDl_L | .992 |
| Thal_MDl_L | Thal_MDl_R | .992 |
| Pallidum_R | Thal_PuM_R | .991 |
| Pallidum_R | Thal_PuM_L | .985 |
| Thal_PuM_R | Thal_PuL_R | .977 |
| Pallidum_L | Thal_MDl_L | .968 |
| Thal_LP_R | Thal_PuA_L | .954 |
| Cingulate_Post_R | Thal_AV_R | .949 |
| Cingulate_Post_R | Thal_PuM_R | .949 |
| Thal_LP_R | Thal_PuA_R | .949 |
| Thal_VL_R | Thal_MDm_L | .949 |
| Thal_VL_R | Thal_MDl_R | .949 |
| Thal_MDm_L | LC_L | .949 |
| Pallidum_R | Thal_PuA_L | .947 |
| Thal_VA_R | Thal_PuM_L | .947 |
| ParaHippocampal_L | Thal_LP_R | .945 |
| Thal_AV_R | Thal_VL_R | .939 |
| Thal_VPL_L | Thal_PuA_R | .939 |
| Thal_MDl_L | Thal_PuM_L | .939 |
| Thal_IL_L | Thal_PuM_R | .938 |
| Pallidum_L | Pallidum_R | .937 |
| Thal_LP_R | Thal_MDl_L | .933 |
| Thal_MDm_L | Thal_PuM_L | .931 |
| Thal_MDm_L | Thal_PuM_R | .931 |
| Pallidum_R | Thal_MDm_L | .919 |
| Cingulate_Post_R | Pallidum_L | .917 |
| Thal_PuL_R | N_Acc_L | .914 |
| Thal_MDl_R | Thal_LGN_L | .910 |
| Thal_VPL_L | Thal_PuM_L | .907 |
| Thal_AV_R | Thal_VPL_L | .905 |
| Thal_AV_R | Thal_PuM_L | .905 |

| **Supplementary Table 14.** Nodal degree and nodal strength of brain regions having highly robust connections identified by CPM (selection frequency≥.9; combined model). | | |
| --- | --- | --- |
| **Label** | **Nodal Degree** | **Nodal Strength** |
| Thal_PuA_L | 15 | 14.740 |
| Thal_MDm_L | 15 | 14.540 |
| Thal_MDl_R | 12 | 11.880 |
| Thal_PuM_R | 10 | 9.740 |
| Thal_LP_R | 10 | 9.580 |
| Thal_AV_R | 9 | 8.740 |
| Thal_PuM_L | 8 | 7.880 |
| Thal_MDm_R | 7 | 6.820 |
| Pallidum_R | 7 | 6.800 |
| Thal_VL_L | 6 | 5.940 |
| Thal_MDl_L | 6 | 5.920 |
| Thal_VPL_L | 6 | 5.920 |
| Thal_VL_R | 6 | 5.900 |
| Thal_PuA_R | 6 | 5.720 |
| Cingulate_Post_R | 4 | 3.840 |
| Pallidum_L | 4 | 3.820 |
| Thal_LGN_L | 3 | 2.880 |
| Thal_VA_L | 3 | 2.880 |
| Thal_VA_R | 3 | 2.860 |
| Thal_PuL_R | 2 | 1.920 |
| Cingulate_Post_L | 2 | 1.860 |
| Frontal_Med_Orb_L | 1 | 1.000 |
| Thal_PuI_R | 1 | 1.00 |
| Thal_AV_L | 1 | 1.00 |
| Thal_VPL_R | 1 | .980 |
| LC_L | 1 | .980 |
| ACC_pre_L | 1 | .940 |
| Vermis_9 | 1 | .940 |
| ParaHippocampal_L | 1 | .940 |
| Thal_PuL_L | 1 | .920 |
| VTA_L | 1 | .920 |
| Cerebellum_Crus1_R | 1 | .900 |
| Putamen_L | 1 | .900 |
| **Note:** Nodal strength was computed from the selection frequency of connections linking the corresponding brain regions. | | |

| **Supplementary Table 15.** Robust connections identified by CPM (selection frequency≥.9; combined model). | | |
| --- | --- | --- |
| **Region A** | **Region B** | **Selection Frequency** |
| Frontal_Med_Orb_L | Pallidum_L | 1.000 |
| Cingulate_Post_R | Thal_VL_L | 1.000 |
| Pallidum_R | Thal_LGN_L | 1.000 |
| Pallidum_R | Thal_PuI_R | 1.000 |
| Pallidum_R | Thal_PuM_R | 1.000 |
| Thal_AV_L | Thal_PuM_R | 1.000 |
| Thal_AV_R | Thal_VL_R | 1.000 |
| Thal_AV_R | Thal_MDm_L | 1.000 |
| Thal_AV_R | Thal_MDl_R | 1.000 |
| Thal_LP_R | Thal_MDm_L | 1.000 |
| Thal_LP_R | Thal_MDl_R | 1.000 |
| Thal_LP_R | Thal_PuA_L | 1.000 |
| Thal_VA_L | Thal_PuA_L | 1.000 |
| Thal_VA_R | Thal_PuM_L | 1.000 |
| Thal_VL_L | Thal_MDm_L | 1.000 |
| Thal_VL_L | Thal_MDm_R | 1.000 |
| Thal_VL_L | Thal_PuA_L | 1.000 |
| Thal_VL_R | Thal_MDl_R | 1.000 |
| Thal_VL_R | Thal_PuM_L | 1.000 |
| Thal_VL_R | Thal_PuA_L | 1.000 |
| Thal_VPL_L | Thal_MDm_L | 1.000 |
| Thal_VPL_L | Thal_MDm_R | 1.000 |
| Thal_VPL_L | Thal_MDl_R | 1.000 |
| Thal_MDm_L | Thal_MDm_R | 1.000 |
| Thal_MDm_L | Thal_MDl_R | 1.000 |
| Thal_MDm_L | Thal_PuA_L | 1.000 |
| Thal_MDm_R | Thal_MDl_L | 1.000 |
| Thal_MDm_R | Thal_PuA_L | 1.000 |
| Thal_MDl_L | Thal_MDl_R | 1.000 |
| Thal_MDl_L | Thal_PuM_L | 1.000 |
| Thal_MDl_L | Thal_PuA_L | 1.000 |
| Thal_MDl_R | Thal_PuM_L | 1.000 |
| Thal_MDl_R | Thal_PuM_R | 1.000 |
| Thal_MDl_R | Thal_PuA_L | 1.000 |
| Thal_PuM_R | Thal_PuA_L | 1.000 |
| Thal_PuA_L | Thal_PuA_R | 1.000 |
| Cingulate_Post_R | Thal_PuM_R | .980 |
| Thal_AV_R | Thal_VA_L | .980 |
| Thal_AV_R | Thal_PuM_L | .980 |
| Thal_AV_R | Thal_PuA_L | .980 |
| Thal_LP_R | Thal_VPL_R | .980 |
| Thal_LP_R | Thal_MDl_L | .980 |
| Thal_VL_L | Thal_MDl_R | .980 |
| Thal_VL_R | Thal_MDm_L | .980 |
| Thal_VPL_L | Thal_PuM_R | .980 |
| Thal_VPL_L | Thal_PuA_R | .980 |
| Thal_MDm_L | Thal_PuM_L | .980 |
| Thal_MDm_L | LC_L | .980 |
| Thal_PuM_R | Thal_PuL_R | .980 |
| Pallidum_L | Pallidum_R | .960 |
| Pallidum_R | Thal_PuM_L | .960 |
| Pallidum_R | Thal_PuA_L | .960 |
| Thal_AV_R | Thal_VPL_L | .960 |
| Thal_VA_R | Thal_MDm_L | .960 |
| Thal_VL_L | Thal_PuM_R | .960 |
| Thal_MDl_R | Thal_PuA_R | .960 |
| Thal_PuM_L | Thal_PuA_L | .960 |
| Cingulate_Post_L | Thal_PuA_L | .940 |
| Cingulate_Post_R | Pallidum_L | .940 |
| ParaHippocampal_L | Thal_LP_R | .940 |
| Vermis_9 | Thal_LGN_L | .940 |
| Thal_LP_R | Thal_PuA_R | .940 |
| Thal_LP_R | ACC_pre_L | .940 |
| Thal_MDl_L | Thal_PuM_R | .940 |
| Thal_MDl_R | Thal_LGN_L | .940 |
| Thal_PuA_R | Thal_PuL_R | .940 |
| Cingulate_Post_L | VTA_L | .920 |
| Cingulate_Post_R | Thal_AV_R | .920 |
| Pallidum_L | Thal_VL_R | .920 |
| Pallidum_R | Thal_MDm_L | .920 |
| Thal_AV_R | Thal_MDm_R | .920 |
| Thal_MDm_L | Thal_PuL_L | .920 |
| Putamen_L | Thal_LP_R | .900 |
| Cerebellum_Crus1_R | Thal_PuA_L | .900 |
| Thal_LP_R | Thal_MDm_R | .900 |
| Thal_VA_L | Thal_VA_R | .900 |
| Thal_MDm_L | Thal_PuM_R | .900 |
| Thal_MDm_L | Thal_PuA_R | .900 |

### **Supplementary Figures**

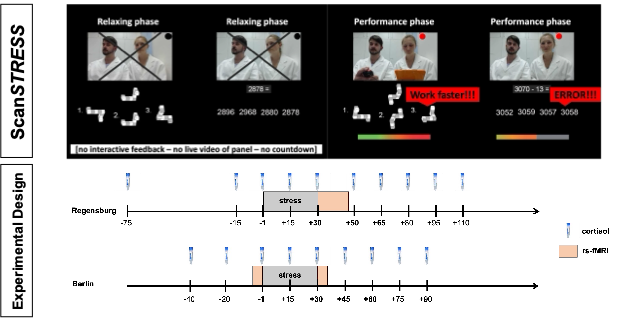

**Supplementary Figure 1.** Illustration of the Scan*STRESS* paradigm, including repeated collection of salivary cortisol and rs‑fMRI images. Adapted from Henze et al. (2025) and Serin et al. (2026). The total duration of the rs‑fMRI acquisition was 18 minutes in Regensburg and 5.18 minutes in Berlin.

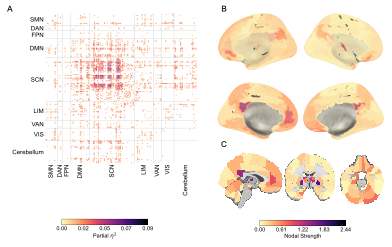
**Supplementary Figure 2.** The NBS sensitivity analysis at (*t* < -2) revealed a significant subnetwork (1564 connections among 164 regions) negatively associated with cortisol increase. Positive associations (*t* > 2) did not reach significance. **A.** The heatmap indicates the intra‑ and inter‑connections, grouped by the assigned canonical functional networks (see Methods for details) and colored according to their effect size ($\eta_{partial}^{2}$). Connections are colored based on their effect size ($\eta_{partial}^{2}$). SMN: Somatomotor Network; DAN: Dorsal Attention Network; FPN: Frontoparietal Network; DMN: Default Mode Network; SCN: Subcortical Network; LIM: Limbic Network; VAN: Ventral Attention Network; VIS: Visual Network. **B.** and **C.** Nodal strength of brain regions found in the significant subnetwork is displayed on the brain surface (fsaverage) and in MNI volumetric space. The details of the nodal strength computation were provided in the Supplementary Methods.

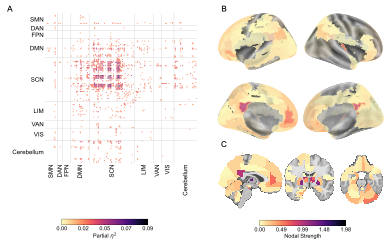
**Supplementary Figure 3.** The NBS sensitivity analysis at (*t* < -2.5) revealed a significant subnetwork (714 connections among 132 regions) negatively associated with cortisol increase. Positive associations (*t* > 2.5) did not reach significance. **A.** The heatmap indicates the intra‑ and inter‑connections, grouped by the assigned canonical functional networks (see Methods for details) and colored according to their effect size ($\eta_{partial}^{2}$). Connections are colored based on their effect size ($\eta_{partial}^{2}$). SMN: Somatomotor Network; DAN: Dorsal Attention Network; FPN: Frontoparietal Network; DMN: Default Mode Network; SCN: Subcortical Network; LIM: Limbic Network; VAN: Ventral Attention Network; VIS: Visual Network. **B.** and **C.** Nodal strength of brain regions found in the significant subnetwork is displayed on the brain surface (fsaverage) and in MNI volumetric space. The details of the nodal strength computation were provided in the Supplementary Methods.

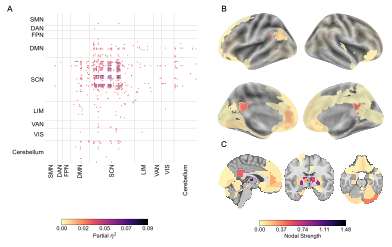
**Supplementary Figure 4.** The NBS sensitivity analysis at (*t* < -3) revealed a significant subnetwork (311 connections among 91 regions) negatively associated with cortisol increase. Positive associations (*t* > 3) did not reach significance. **A.** The heatmap indicates the intra‑ and inter‑connections, grouped by the assigned canonical functional networks (see Methods for details) and colored according to their effect size ($\eta_{partial}^{2}$). Connections are colored based on their effect size ($\eta_{partial}^{2}$). SMN: Somatomotor Network; DAN: Dorsal Attention Network; FPN: Frontoparietal Network; DMN: Default Mode Network; SCN: Subcortical Network; LIM: Limbic Network; VAN: Ventral Attention Network; VIS: Visual Network. **B.** and **C.** Nodal strength of brain regions found in the significant subnetwork is displayed on the brain surface (fsaverage) and in MNI volumetric space. The details of the nodal strength computation were provided in the Supplementary Methods.

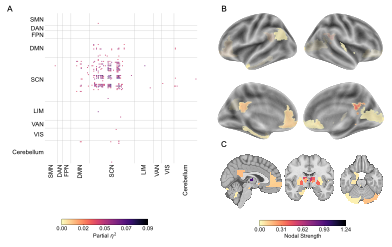
**Supplementary Figure 5.** The NBS sensitivity analysis at (*t* < -3.5) revealed a significant subnetwork (138 connections among 48 regions) negatively associated with cortisol increase. Positive associations (*t* > 3.5) did not reach significance. **A.** The heatmap indicates the intra‑ and inter‑connections, grouped by the assigned canonical functional networks (see Methods for details) and colored according to their effect size ($\eta_{partial}^{2}$). Connections are colored based on their effect size ($\eta_{partial}^{2}$). SMN: Somatomotor Network; DAN: Dorsal Attention Network; FPN: Frontoparietal Network; DMN: Default Mode Network; SCN: Subcortical Network; LIM: Limbic Network; VAN: Ventral Attention Network; VIS: Visual Network. **B.** and **C.** Nodal strength of brain regions found in the significant subnetwork is displayed on the brain surface (fsaverage) and in MNI volumetric space. The details of the nodal strength computation were provided in the Supplementary Methods.

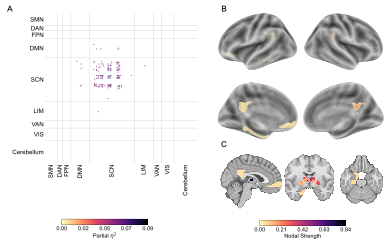
**Supplementary Figure 6.** The NBS sensitivity analysis at (*t* < -4) revealed a significant subnetwork (67 connections among 29 regions) negatively associated with cortisol increase. Positive associations (*t* > 4) did not reach significance. **A.** The heatmap indicates the intra‑ and inter‑connections, grouped by the assigned canonical functional networks (see Methods for details) and colored according to their effect size ($\eta_{partial}^{2}$). Connections are colored based on their effect size ($\eta_{partial}^{2}$). SMN: Somatomotor Network; DAN: Dorsal Attention Network; FPN: Frontoparietal Network; DMN: Default Mode Network; SCN: Subcortical Network; LIM: Limbic Network; VAN: Ventral Attention Network; VIS: Visual Network. **B.** and **C.** Nodal strength of brain regions found in the significant subnetwork is displayed on the brain surface (fsaverage) and in MNI volumetric space. The details of the nodal strength computation were provided in the Supplementary Methods.

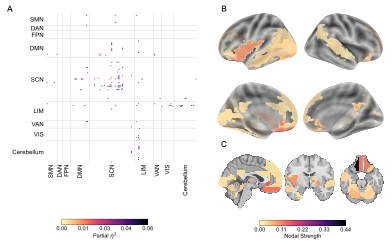
**Supplementary Figure 7.** Results of sex‑specific NBS analyses (male contrast): an uncorrected subnetwork (65 connections among 53 regions) was negatively associated with cortisol increase in males (*t* < –3.1). It did not survive FDR correction, nor did its positive associations (*t* > 3.1) (Supplementary Table 8). **A.** The heatmap indicates the intra‑ and inter‑connections, grouped by the assigned canonical functional networks (see Methods for details) and colored according to their effect size ($\eta_{partial}^{2}$). Connections are colored based on their effect size ($\eta_{partial}^{2}$). SMN: Somatomotor Network; DAN: Dorsal Attention Network; FPN: Frontoparietal Network; DMN: Default Mode Network; SCN: Subcortical Network; LIM: Limbic Network; VAN: Ventral Attention Network; VIS: Visual Network. **B.** and **C.** Nodal strength of brain regions found in the significant subnetwork is displayed on the brain surface (fsaverage) and in MNI volumetric space. The details of the nodal strength computation were provided in the Supplementary Methods.

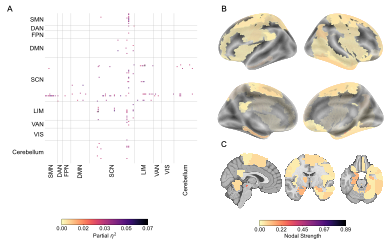
**Supplementary Figure 8.** Results of sex‑specific NBS analyses (male > female): an uncorrected subnetwork (70 connections among 57 regions) was negatively associated with cortisol increase in females (*t* < –3.1). It did not survive FDR correction (Supplementary Table 8). **A.** The heatmap indicates the intra‑ and inter‑connections, grouped by the assigned canonical functional networks (see Methods for details) and colored according to their effect size ($\eta_{partial}^{2}$). Connections are colored based on their effect size ($\eta_{partial}^{2}$). SMN: Somatomotor Network; DAN: Dorsal Attention Network; FPN: Frontoparietal Network; DMN: Default Mode Network; SCN: Subcortical Network; LIM: Limbic Network; VAN: Ventral Attention Network; VIS: Visual Network. **B.** and **C.** Nodal strength of brain regions found in the significant subnetwork is displayed on the brain surface (fsaverage) and in MNI volumetric space. The details of the nodal strength computation were provided in the Supplementary Methods.

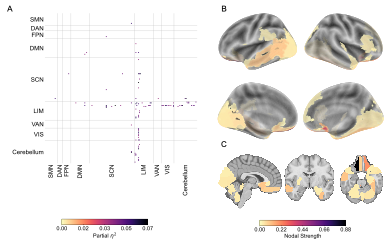
**Supplementary Figure 9.** Results of sex‑specific NBS analyses (female > male): an uncorrected subnetwork (51 connections among 48 regions) was negatively associated with cortisol increase in females (*t* < –3.1). It did not survive FDR correction (Supplementary Table 8). **A.** The heatmap indicates the intra‑ and inter‑connections, grouped by the assigned canonical functional networks (see Methods for details) and colored according to their effect size ($\eta_{partial}^{2}$). Connections are colored based on their effect size ($\eta_{partial}^{2}$). SMN: Somatomotor Network; DAN: Dorsal Attention Network; FPN: Frontoparietal Network; DMN: Default Mode Network; SCN: Subcortical Network; LIM: Limbic Network; VAN: Ventral Attention Network; VIS: Visual Network. **B.** and **C.** Nodal strength of brain regions found in the significant subnetwork is displayed on the brain surface (fsaverage) and in MNI volumetric space. The details of the nodal strength computation were provided in the Supplementary Methods.

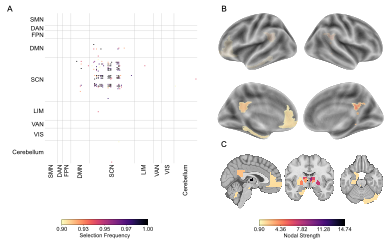
**Supplementary Figure 10.** Stable connections (78 connections among 33 regions), predicting cortisol increase from using CPM (combined network; *r* = .173, *p_FDR_* = .009). A selection frequency threshold of .90 was used to visualize the stable connections selected across cross-validation folds. **A.** The heatmap indicates the intra‑ and inter‑connections, grouped by the assigned canonical functional networks (see Methods for details) and colored according to their selection frequency values. SMN: Somatomotor Network; DAN: Dorsal Attention Network; FPN: Frontoparietal Network; DMN: Default Mode Network; SCN: Subcortical Network; LIM: Limbic Network; VAN: Ventral Attention Network; VIS: Visual Network. **B.** and **C.** Nodal strengths of stable brain regions are displayed on the brain surface (fsaverage) and in MNI volumetric space. The details of the nodal strength computation were provided in the Supplementary Methods.

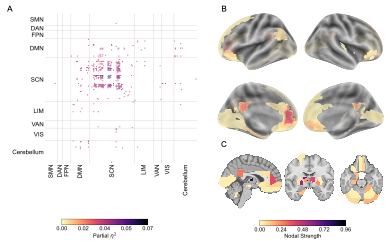

**Supplementary Figure 11.** The NBS analysis uses truncated rs-fMRI data from the Regensburg site. To match the acquisition duration from the Berlin and Regensburg sites, only a limited number of scanning volumes were used to generate connectivity matrices in Regensburg. A significant subnetwork (192 connections among 71 regions) negatively associated with cortisol increase was revealed by NBS analysis (*t*<-3.1). Positive associations (*t*>3.1) did not reach significance. **A.** The heatmap indicates the intra‑ and inter‑connections, grouped by the assigned canonical functional networks (see Methods for details) and colored according to their effect size ($\eta_{partial}^{2}$). Connections are colored based on their effect size ($\eta_{partial}^{2}$). SMN: Somatomotor Network; DAN: Dorsal Attention Network; FPN: Frontoparietal Network; DMN: Default Mode Network; SCN: Subcortical Network; LIM: Limbic Network; VAN: Ventral Attention Network; VIS: Visual Network. **B.** and **C.** Nodal strength of brain regions found in the significant subnetwork is displayed on the brain surface (fsaverage) and in Montréal Neurological Institute (MNI) volumetric space. The details of the nodal strength computation are provided in the Supplementary Methods.
